# Exploration and exploitation are flexibly balanced during local search in flies

**DOI:** 10.1101/2024.06.26.600764

**Authors:** Dennis Goldschmidt, Yipei Guo, Shivam S Chitnis, Christina Christoforou, Dan Turner-Evans, Carlos Ribeiro, Ann M Hermundstad, Vivek Jayaraman, Hannah Haberkern

## Abstract

After finding food, a foraging animal must decide whether to continue feeding, or to explore the environment for potentially better options. One strategy to negotiate this tradeoff is to perform local searches around the food but repeatedly return to feed. We studied this behavior in flies and used genetic tools to uncover the underlying mechanisms. Over time, flies gradually expand their search, shifting from primarily exploiting food sources to exploring the environment, a change that is likely driven by increases in satiety. We found that flies’ search patterns preserve these dynamics even as the overall scale of the search is modulated by starvation-induced changes in metabolic state. In contrast, search induced by optogenetic activation of sugar sensing neurons does not show these dynamics. We asked what navigational strategies underlie local search. Using a generative model, we found that a change in locomotor pattern after food consumption could account for repeated returns to the food, but failed to capture relatively direct, long return trajectories. Alternative strategies, such as path integration or sensory taxis could allow flies to return from larger distances. We tested this by individually silencing the fly’s head direction system, olfaction and hygrosensation, and found that the only substantial effect was from perturbing hygrosensation, which reduced the number of long exploratory trips. Our study illustrates that local search is composed of multiple behavioral features that evolve over time based on both internal and external factors, providing a path towards uncovering the underlying neural mechanisms.

## Introduction

A sound foraging strategy involves not just finding food, but also making appropriate decisions afterwards to achieve nutrient homeostasis. Should the animal stop at the food source? How long should it stay and feed? Should it search to identify other, potentially richer food sources? How far should it search before it returns to the known resource? Many species, ranging from insects (Bell, 1985, 1990; Visser, 1988) to mammals (Bailey et al., 2019; Simpkins et al., 2001; Thiele and Winter, 2005), including humans (Jang et al., 2019; Pacheco-Cobos et al., 2019), rely on an adaptive local search around food for survival. This ubiquitous foraging behavior combines exploratory movements around the discovered food with exploitative bouts of feeding during returns.

Classical work from Vincent Dethier (Dethier, 1957) described hungry blowflies increasing their turning after feeding from a small drop of sugar. The shape and intensity of these searches were modulated by the sugar concentration, as well as how long the fly was starved (Bell et al., 1985; Nelson, 1977). Since the tiny food spot used in these experiments was likely consumed during the first encounter, it was argued that the search could be a means to look for the lost resource. Alternatively, searching within the vicinity of a found resource could also be a tactic to locate a different resource (Bell, 1985). This behavior was thought to reflect a prior on the distribution of food resources: if food was inhomogeneously distributed in patches in the environment, a local search around a previously identified food source was likely to reveal further resources. Potentially, these other food sources could provide more valuable nutrients for the animal than the first encountered spot. If the initial food spot is not consumed during the first encounter, local search can therefore be interpreted as balancing exploration of the environment with exploitation of a known resource, based on an innate expectation of the patchy distribution of resources in nature (Hills et al., 2015).

Internal states shape foraging behavior by influencing both food choice and foraging strategies (Münch et al., 2020). Hunger and satiety signals jointly support nutrient homeostasis: Nutrient deprivation induces nutrient-specific appetites or hunger, which drives the animal to seek out and consume those nutrients that are lacking (Ribeiro and Dickson, 2010; Steck et al., 2018), while satiety signals indicate when physiological needs are met to prevent overconsumption (Malita et al., 2022; Sun et al., 2017). Many studies have shown how metabolic hunger and satiety states shape behavior in the context of feeding regulation and nutrient food choice (Itskov et al., 2014; Ribeiro and Dickson, 2010; Yapici et al., 2016). Mechanistically, these internal states impinge on neuronal circuits at different levels in brain processing (Münch et al., 2020), ranging from gain modulation in sensory systems (Inagaki et al., 2014; Marella et al., 2012; Munch et al., 2022; Steck et al., 2018) to modulating motor systems that control feeding (Munch et al., 2022; Pool et al., 2014). Nutrient-specific appetites and satiety signals not only affect feeding but also foraging decisions. Exploitative and exploratory strategies are modulated in amino-acid deprived flies, which reduces global exploration and increases local exploration of protein-rich food sources (Corrales-Carvajal et al., 2016). While first steps towards identifying neuronal substrates for regulating specific decision parameters during foraging have been made (Goldschmidt et al., 2023), the neural circuit logic and the dynamic interplay between hunger and satiety states in regulating foraging behavior remain less well understood.

Previous descriptions of long-timescale behaviors, such as local search for food, have lacked accurate, high-resolution quantifications of movement. Recent technological advances have enabled quantification of increasingly naturalistic behaviors at high spatial and temporal resolution (Alisch et al., 2018; Barlow et al., 2022; Bath et al., 2014; Branson et al., 2009; Mathis et al., 2018). Studies in flies and bees have used these advances to gain insights into navigational aspects of local search in insects, exploring primarily how animals return to a known food source (Brockmann et al., 2018; Corrales-Carvajal et al., 2016; Kim and Dickinson, 2017; Murata et al., 2017; Shakeel and Brockmann, 2023; Titova et al., 2023). Later studies have substituted the real food with optogenetic activation of various sensory and brain neurons, to simplify the assay and identify potential neuronal elements involved in eliciting search (Corfas et al., 2019; Haberkern et al., 2019). To further reduce complexity, more recent studies have restricted search to a single dimension (Behbahani et al., 2021; Corfas et al., 2019). However, these simplifications of substituting real food with optogenetic stimulation and restricting the animal’s movement may mask how the tradeoff between exploration and exploitation is dynamically managed during local search and how food consumption affects the behavior.

In this study, we quantitatively analyze the local search patterns of fruit flies in an open field assay with real food. By tracking the movement paths at high temporal and spatial resolution, we dissect the fine-scale maneuvers that constitute a local search and evaluate how these patterns change over time and with changing environmental and physiological conditions. We explore the balance between the exploration and exploitation phases of local search, investigating how shortterm food encounters and long-term starvation states modulate these behaviors. Our results expand our understanding of foraging theory and apply a quantitative lens to the complex decision-making processes underlying animal behavior in natural settings

## Results

We studied local search behavior over 45 minutes in 24 hours (h) starved, mated, female flies (*Drosophila melanogaster*) in a large, dark arena with a 5 µl food spot (**Fig. 1A**, Methods). Upon finding the food, flies drastically changed their movement pattern to generate a food-centered search (Bell, 1985; Corrales-Carvajal et al., 2016; Dethier, 1957; Kim and Dickinson, 2017; Murata et al., 2017). We sought to investigate the spatiotemporal structure of this search, relating exploratory behavior to food encounters. We therefore segmented trajectories into segments defined by flies’ movements, their contact with the food and the arena border (**Fig. 1B**, Methods):

**Figure 1.**
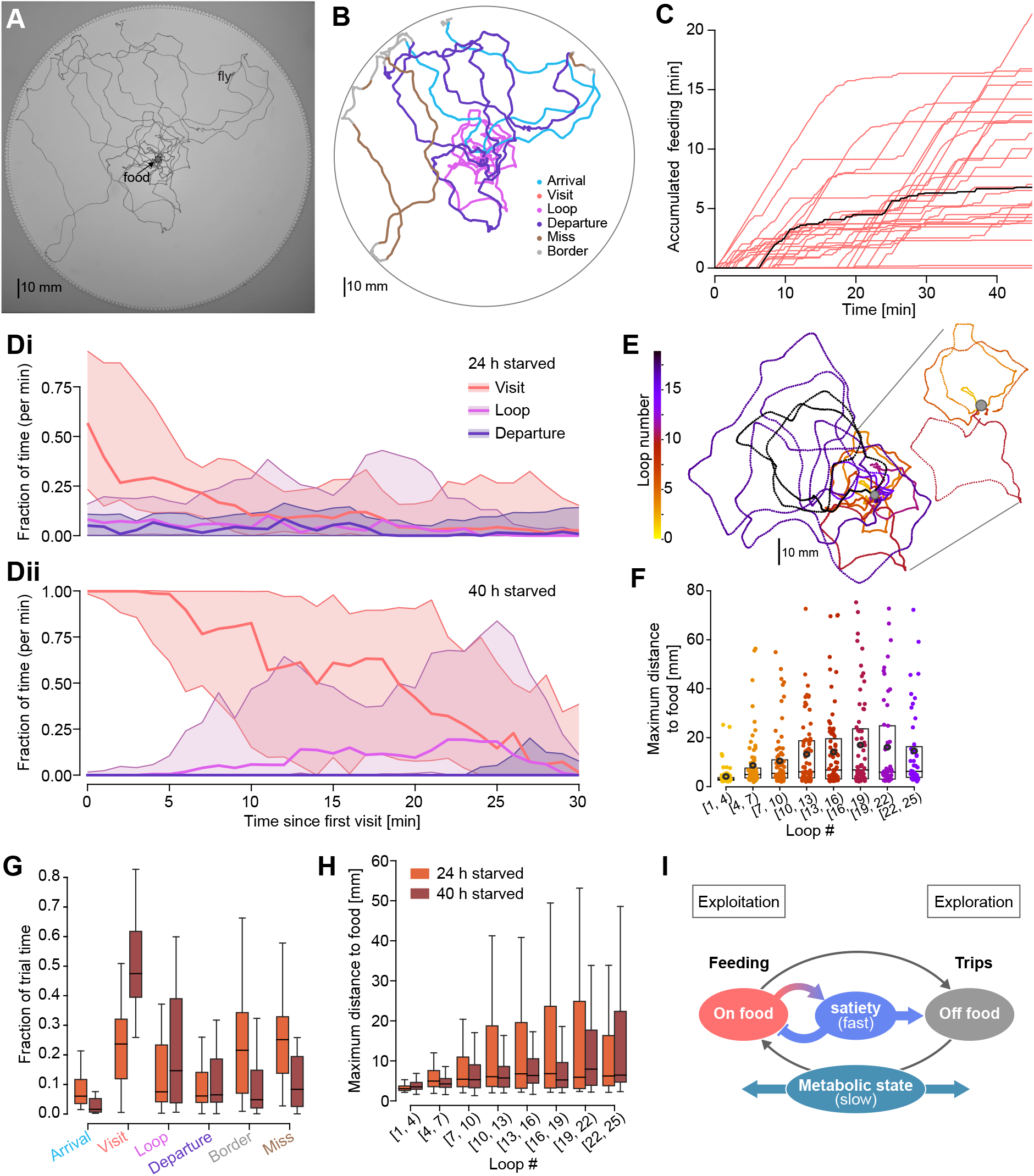
Local search for food is structured in space and time. A: Walking trajectory of a 24 h starved fly (fraction of the 45 min recording). An image frame from the recorded video is shown for reference. The food spot (0.125 M sucrose in agarose) was positioned in the center of the arena. B: Same trajectory as in A color coded by the assigned segment type. C: Accumulated feeding time over the course of the trial for each fly in the 24 h starved sucrose food group. Each line corresponds to a different fly. The fly shown in A, B and E is highlighted in black. D: Time series showing the average fraction per minute that flies spent a given behavioral segment (visits, loops, departures). The fraction was computed for each fly using a forward-facing, 5 min wide rolling time window. Time series were aligned to the time point when flies made their first visit. Median (line) and interquartile range (shaded region) are shown 24 h starved flies (*n* = 28) in Di and 40 h starved flies (*n* = 29) in Dii. E: Loop sections of the same trajectory as shown in A & B with individual loops color coded according to their temporal order. Left shows a zoom-in on early loops. F: Strip plot of the maximum distance from the food that was reached in each loop with loops binned according to their temporal order (24 h starved flies with sucrose food, n=27 flies made loops and were included). Black circles mark the mean, the simplified boxplots the median and interquartile range. G: Boxplot showing the fraction of the trial time that flies spent in the different segments (see B and Methods for segment definition). The two experimental groups are color coded: 24 h starved (orange, *n* = 28), and 40 h starved (dark red, *n* = 29). H: Boxplot of the maximum distance from the food that was reached in each loop with loops binned according to their temporal order (see also F), comparing the 24 h and 40 h starved flies. I: Schematic of the conceptual framework applied to local search behavior in this study. See also Supplemental Figure 1.

- “Visits”: segments in which flies were on the food spot. We assumed that micromovements during visits are a reliable indicator of feeding bouts (Corrales-Carvajal et al., 2016).
- “Borders”: segments when flies remained close to the arena border.
- “Departures”: segments after a food visit, which ended at the border.
- “Arrivals”: segments after a border segment, which ended on the food.
- “Misses”: segments which left the border and returned to the border.
- “Loops”: segments in which flies left the food and returned to it without any border contact.

By distinguishing loops from search trajectories in which flies are close to the border, we could selectively analyze trajectory segments where the fly was likely engaging in a local search.

### Local search balances exploration with exploitation based on the fly’s metabolic state

We used segmented search trajectories to investigate how the dynamics of flies’ feeding and exploration developed over time. We found that flies discovered the food and started feeding at different times during the trial, but the dynamics of their feeding bouts were similar: flies tended to eat more immediately after finding the food, and feeding rates slowed down later as they became satiated (**Fig. 1C**). To analyze the temporal dynamics of flies’ search pattern, we aligned their segmented trajectories to the start of their first food visit and computed the fraction of time they spent in a specific behavioral segment (**Fig. 1Di**). Flies initially spent a large part of their time on the food spot (**Fig. 1Di**, “visits”). However, after a few minutes, they gradually devoted more time to searching around the food (**Fig. 1Di**, “loops”) or transitioned to exploration further away (**Fig. 1Di**, “departures”). For search loops, we observed additional spatiotemporal structure: loops gradually increased in size, reaching greater distances from the food spot over subsequent visits (**Fig. 1E**). This scaling of the search pattern was also visible at the population level (**Fig. 1F**, see also **Fig**. **S1D**), although it primarily manifested in an increased variance of loop sizes, rather than as a large increase in the mean loop size. Like previous re-ports (Corrales-Carvajal et al., 2016), our data suggests that flies search farther as they get satiated (**Fig. 1C, F**). To further test this, we compared the behavior of 24 h versus 40 h starved flies. We observed that 40 h starved flies found the food more quickly, stayed longer on the food, and fed more (**Fig. S1A-C**). Longer starved flies also spent more time on local exploration (**Fig. 1Dii, G**, see loops and visits) and expanded their search radii more slowly (**Fig. 1H, Fig. S1D**). This is consistent with previous reports that metabolic states, including those induced by prolonged starvation, have profound effects on feeding and foraging behavior (Bell, 1985; Corrales-Carvajal et al., 2016; Dethier, 1957; Yapici et al., 2016). Despite these differences in the overall search behavior of 24 h and 40 h starved flies, we noted that the temporal structure with longer feeding bouts and local exploration in the beginning of the trial, and increasing spatial exploration later in the trial, is preserved. Thus, we hypothesize that two internal, dynamic variables differentially shape the spatiotemporal structure of local search: a rapidly changing one, possibly based on satiety from recent feeding, drives changes in how the fly explores the surrounding environment (see also Corrales-Carvajal et al. (2016)) and a slowly changing one based on previous food deprivation affects the overall scale (**Fig. 1I**). Next, we aim to tease out potential mechanisms underlying these dynamics.

### Metabolic states modulate different aspects of exploitative strategies to achieve nutrient homeostasis

To show how metabolic states affected flies’ exploitative strategies, we quantified how flies interacted with the food spot after 24 and 40 h of starvation (**Fig. 2A**). The increase in feeding for longer starved flies (**Fig. 1G, Fig. 2B**) could come from flies staying longer on the food, visiting the spot more frequently, or both. To distinguish between these possibilities, we examined both the duration and the number of visits across 24 h and 40 h starved flies (**Fig. 2C**, Methods). Evaluating isoclines of total feeding times, which are the product of the number and duration of visits (Corrales-Carvajal et al., 2016), we found that individual animals adopted quite different strategies with some flies increasing the duration of their visits, while others increased the frequency. Overall, flies that were starved longer (40 h) exhibited a significantly longer mean duration of visits (**Fig. S2A**), but the total number of visits across the population was not significantly changed by starvation (**Fig. 2D**). Nutrient deprivation has been shown to modulate both the number of encounters that flies have with food spots and the probability that they engage with and feed on an encountered spot (Corrales-Carvajal et al., 2016; Goldschmidt et al., 2023). Interestingly, neither of these aspects of flies’ foraging behavior changed with starvation in our paradigm (**Fig. S2B** and **Fig. S2C**, respectively). In summary, our data demonstrates that flies’ state of starvation modulates their foraging and feeding strategies. Flies specifically adapt their feeding behavior by modifying the duration and, to a lesser extent, the frequency of feeding visits, rather than altering encounter rates with food spots.

**Figure 2.**
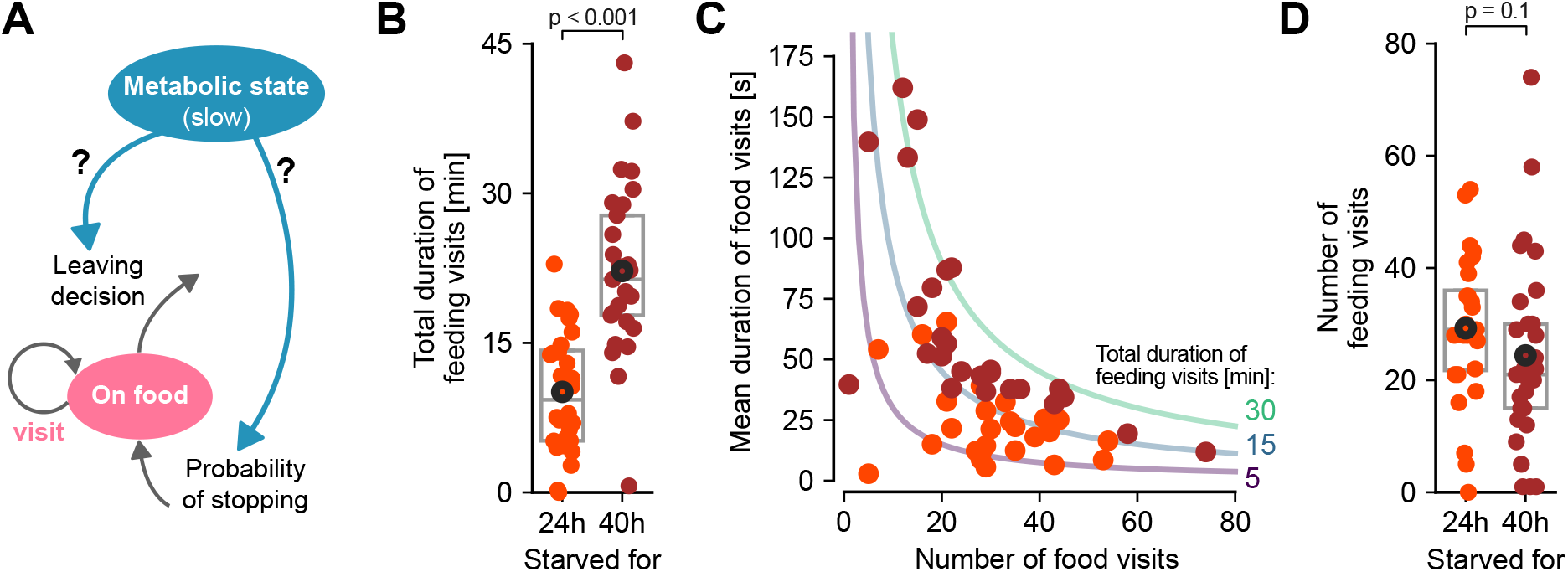
Flies use multiple exploitative strategies to fulfill their metabolic needs. A: Illustration of how the metabolic state (starvation duration) can regulate exploitation through modulating “leaving decisions” and/or the “probability to engage” with a food spot. B: Total duration of feeding visits in minutes (min) for 24 h (orange circles) and 40 h (maroon circles) starved flies. Data points are calculated for individual flies. Boxes indicate the median and interquartile range (IQR, i.e., between 25th and 75th percentile), and circle markers indicate the mean. P-values are obtained by performing Wilcoxon rank-sum tests, *n* = 28 – 29. C: To reach different total durations of feeding visits (purple to green lines), flies may modulate how often to visit (x-axis: number of visits) and/or how long to visit (y-axis: mean duration of feeding visits in min). Data points are calculated for individual, 24 h (orange circles) and 40 h (maroon circles) starved flies. See also Supplemental Figure 2.

### Nutritive and sweet taste cues are important for eliciting and sustaining local search

Once flies encounter a food spot, they use taste and post-ingestive nutritional cues to evaluate the value and quality of the food resource (Burke and Waddell, 2011; Dus et al., 2011; Fujita and Tanimura, 2011; Stafford et al., 2012; Steck et al., 2018). To differentiate between sensory and nutritive value effects on local search, we aimed to separate them by providing food spots with different nutritional value. We tested individual, sucrose-deprived flies with either 100 mM sucrose, 100 mM D-sorbitol, or 100 mM sucralose presented in agarose. D-sorbitol is a sugar alcohol, which does not taste sweet to flies, but which can be metabolized and ensure their survival (Fujita and Tanimura, 2011). In contrast, sucralose is a non-caloric artificial sweetener, which is perceived by flies as sweet but which cannot be metabolized and therefore lacks nutritive value (Abu et al., 2018; Burke and Small, 2015; Park et al., 2017). We observed that the total duration of feeding visits was decreased for sorbitol and sucralose compared to sucrose (**Fig. 3A**). This reduced time on non-sweet and non-nutritive food spots was explained by a higher probability of the fly to leave the spot, indicated by lower mean durations of feeding visits (**Fig. 3B, Fig. S3A**), while the frequency of visits to the different food spots was not significantly different (**Fig. S3A**, **B**). Curiously, feeding times for sorbitol food showed a bimodal distribution (**Fig. 3A**) and those flies that initiated feeding increased their feeding rates during the assay (**Fig. 3C**). This observation could be explained by a mechanism in which flies, which happen to ingest significant amount of the food without taste, experience a positive metabolic feedback signal. This may allow them to overcome the lack of gustatory input and alter their behavior upon the metabolic signal. For sucralose, the amount and rate of feeding visits were both lower than for flies exposed to the sucrose and sorbitol food spots (**Fig. 3A, Fig. S3B**). Likewise, flies exposed to sorbitol and sucralose showed significantly fewer loops compared to those exposed to a sucrose spot (**Fig. S3C**). To further evaluate nutrient-elicited search without taste input, we defined search bouts, in which a fly alternates between feeding and loops without reaching the border (**Fig. S3D**). While search durations were increased in flies exposed to sucrose and sorbitol compared to sucralose (**Fig. 3D**), search distances were comparable between sucrose and sorbitol (**Fig. 3E**). In a separate experiment, we removed both sweet taste and nutritive value from the food spot by providing water with 1% agarose. Flies foraging for water showed significantly lower feeding, both in how long they stay at a food spot as well as in their visit frequency (**Fig. S4**). Taken together, our data suggests that both post-ingestive metabolic and sweet taste cues are necessary to elicit the full search pattern and that post-ingestive cues might influence search behavior on a slower timescale.

**Figure 3.**
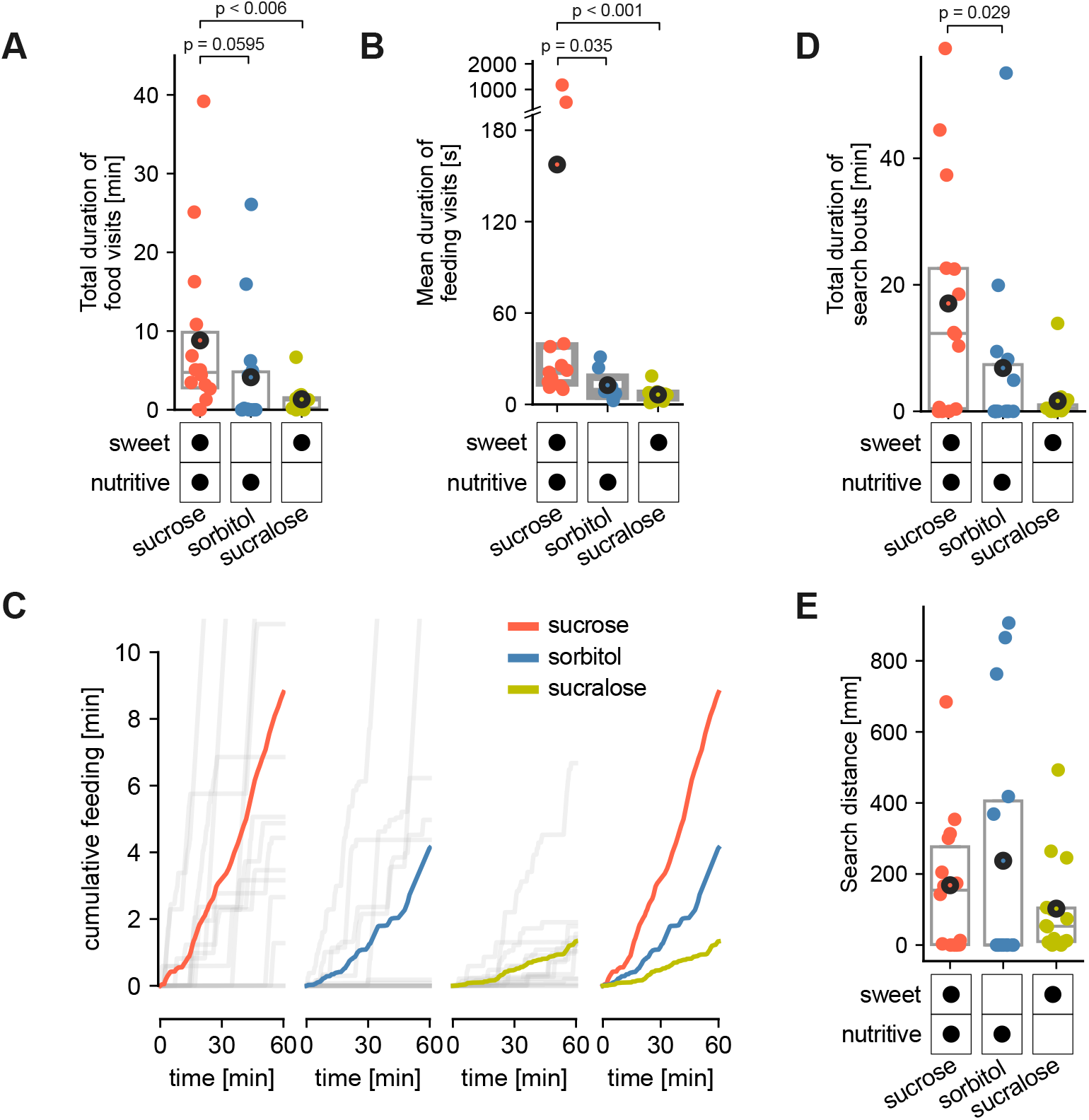
Nutritive and taste cues contribute to eliciting and sustaining search. A: Total duration of feeding visits in minutes (min) for deprived flies presented with 100 mM sucrose (red circles), 100 mM sorbitol (blue circles) or 100 mM sucralose (olive green circles), respectively. Data points are calculated for individual flies. Boxes indicate the median and interquartile range (IQR, i.e., between 25th and 75th percentile), and circle markers indicate the mean. P-values are obtained by performing Wilcoxon rank-sum tests, n = 14. B: To reach different total durations of feeding visits (purple to green lines), flies may modulate how often to visit (x-axis: number of visits) and/or how long to visit (y-axis: mean duration of feeding visits in min). Data points are calculated for individual, 24 h starved flies presented with 100 mM sucrose (red circles), 100 mM sorbitol (blue circles) or 100 mM sucralose (olive green circles), respectively. C: Cumulative feeding in minutes over the course of the trial for each fly (gray lines) as well as the average across flies presented with 100 mM sucrose (red line), 100 mM sorbitol (blue line) or 100 mM sucralose (olive green line). D: Total duration of search bouts in min for deprived flies presented with 100 mM sucrose, 100 mM sorbitol or 100 mM sucralose. E: Search distance in mm for deprived flies presented with 100 mM sucrose, 100 mM sorbitol or 100 mM sucralose. D-E. Data points are calculated for individual flies. Boxes indicate the median and interquartile range (IQR, i.e., between 25th and 75th percentile), and circle markers indicate the mean. P-values are obtained by performing Wilcoxon rank-sum tests, *n* = 1 4. See also Supplemental Figures 3 and 4.

### Optogenetically simulating sweet taste without nutritional value induces searches without temporal scaling

To further explore whether increasing satiety levels are indeed the driving force behind the temporal scaling of the search pattern (**Fig. 4A**), we investigated the spatiotemporal dynamics of search elicited by optogenetic “sweet” stimulation that lacks the nutritive component. We took advantage of a published dataset (Corfas et al., 2019), where starved flies were stimulated for 1 s by optogenetically activating neurons expressing sweet gustatory receptors (Miyamoto and Amrein, 2014; Miyamoto et al., 2012) whenever they reached a restricted area in the center of a walking arena. In this assay, flies responded with a search around the site of stimulation, which we will term “optogenetic search” (Kim and Dickinson, 2017; Titova et al., 2023). To evaluate which features were shared between real food-induced local search and optogenetic search, we segmented and analyzed trajectories from flies that were stimulated by activating neurons targeted by a Gr43a- and Gr5a-Gal4 line (Corfas et al., 2019). As described in the original studies, optogenetic search, like food-induced search, led to extensive looping around the stimulation site (**Fig. 4B-D**), but flies spent more time at the border and often missed the virtual food source (**Fig. 4C**). Further, optogenetic search behavior was generally more variable, with some flies responding strongly (**Fig. 4B**) and others performing just a few loops (**Fig. 4D**). Consistent with the assumption that changes in satiety underly the gradual expansion of the search pattern, we found that loops in the optogenetic search did not show a systematic increase in loop distances over time (**Fig. 4E**, compare also **Fig. 4B** with **Fig. 1E, Fig. S5A**). Also, in contrast to real food-induced searches, optogenetic search showed greater curvature (**Fig. 4B** vs. **Fig. 1E**) and higher tortuosity of long loops (**Fig. 4F, Fig. S5B-E**). Despite, or perhaps because of this altered walking pattern, optogenetically stimulated flies showed return rates to the stimulation site that were comparable to those seen in real food search (**Fig. S5F-G**). Notably, the return rates and scale of the search in the 40 h starved, optogenetically stimulated flies resembled the phenotype of 24 h more than 40 h starved flies in real food search, suggesting that the effect of metabolic state on optogenetic and real food-induced search may differ as well. These findings show that sweet stimuli generated through activation of gustatory neurons alone are insufficient to elicit the full food-induced local search pattern and suggest that additional signals from interacting with or ingesting food are necessary to generate the slow expansion of search over time.

**Figure 4.**
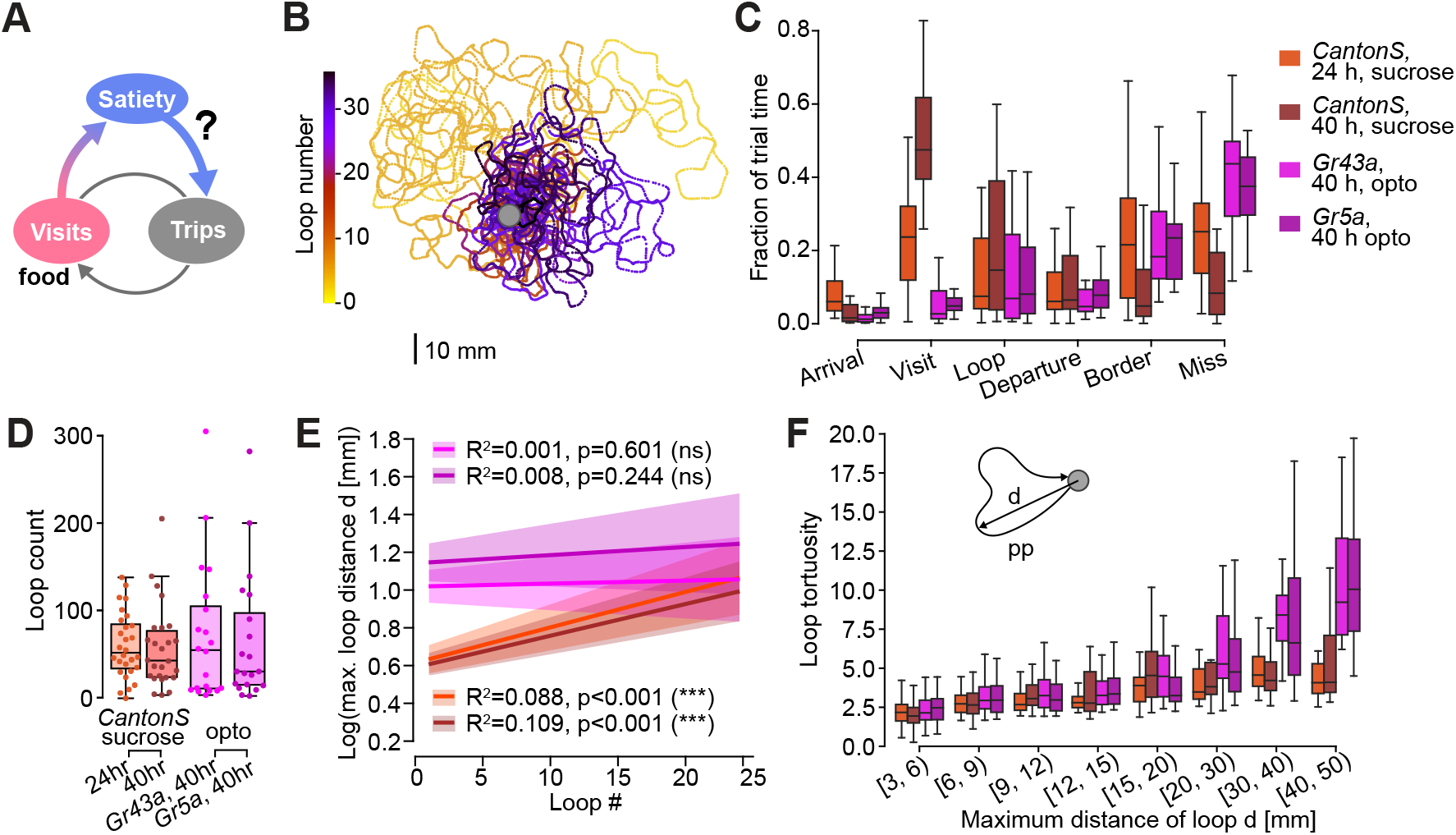
Optogenetic activation of sugar sensing neurons without food ingestion does not induce the full local search pattern. A: Possible interaction of “on food” (visits) and “off food” (trips) behavior via the satiety state. B: Loop sections of a trajectory from a fly after optogenetic stimulation of *Gr43a* neurons. Data from 37. Loops are color coded according to their temporal order as in Figure 1D. C: Boxplot showing the fraction of the trial time that flies spent in the different segments (same as in Fig. 1G). Experimental groups are color coded: wild type with sucrose food, 24 h starved (orange, *n* = 28), and 40 h starved (dark red, *n* = 29); optogenetically stimulated flies from 37 using Gr43a-Gal4 (pink, *n* = 20) and Gr5a-Gal4 (dark purple, *n* = 20). D: Total number of loops per fly for the four experimental groups (color coded as in C). E: Results of ordinary least squares linear regression of the log(maximal radial loop distance [mm]) as a function of the loop number. The log transform was chosen to account for the long-tailed distribution of loop distances. Groups are color coded as in C. The lines indicate the regression line, the shaded regions the 95% confidence intervals. Full parameters: Gr43a>CsChrimson: R^2^=0.001, p=0.6005009258188643 (ns); Gr5a>CsChrimson: R^2^=0.008, p=0.24412469659864977 (ns); 24 h starved *Canton-S*: R^2^=0.088, p=6.270106396335541e-12 (***); 40 h starved *Canton-S*: R^2^=0.109, p=9.417834233499424e-14 (***). F: Ratio of path length [mm] and maximum radial distance from the food [mm] for loops. Note that uneven bin sizes were chosen to account for reduced number of samples for larger distances from the food. See also Supplemental Figure 5.

### Search trips can be divided into two groups based on their duration

To decipher the mechanisms driving search expansion, we closely analyzed the pattern of search loop lengths. We had noted that despite the overall scaling of the search pattern over time flies frequently performed short loops late in the search sequence (**Fig. 1F**). We therefore investigated the distribution of path lengths of individual search trips, that is, all trajectory segments that started and ended at the food, including those that reached the arena border. We found a segregation into two types of trips: long and short ones (**Fig. 5A, B**). To account for fly-specific differences, we normalized the trip path lengths per fly. Indeed, the distribution of normalized, log-transformed trip lengths showed a bimodal shape, which was fit by two gaussians (**Fig. 5B**, see **Fig. S6A-B** for loop-based analysis). We used the intersection point of the two fitted gaussians to classify trips into short and long ones (**Fig. 5B-D**). In 40 h starved flies, the separation into two trip types was less clear (**Fig. S6C-D**), possibly due to the reduced spatial extent of the search pattern (**Fig. 1H**). Given the evidence for two types of trips, we analyzed long and short trips separately to look for possible trip-type specific adaptations that shape the search pattern.

**Figure 5.**
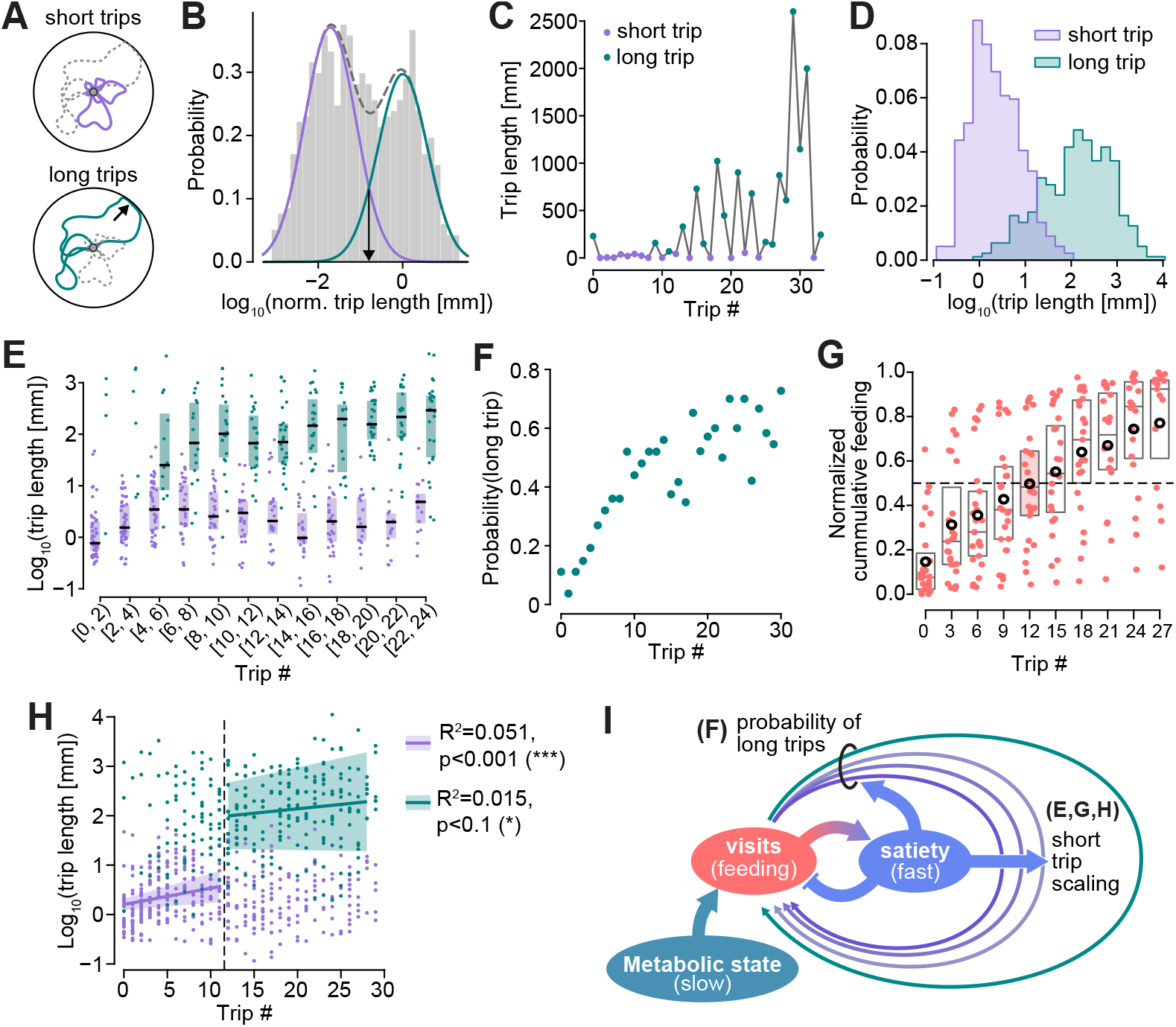
Two types of search trips adapt differentially to generate an expanding search pattern. A: Distinction between short trips (loops) and long trips (long loop segments and excursions composed of departure, border and arrival segments). B-I: Analysis of short and long trips for wild type, 24 h starved flies on sucrose food (*n* = 28). See supplemental data for the 40 h group. B: Bimodal distribution of log-transformed, normalized trip lengths (gray shaded histogram, see methods for details). Trip lengths were normalized per fly by dividing each trip (path) length by the average trip length. The distribution can be approximated by two gaussians corresponding to short trips (purple, mean = −1.71, sd = 0.61, weight = 0.56) and long trips (teal, mean = 0.0, sd = 0.59, weight = 0.44). The intersection point of the two gaussians (black arrow) is used as a threshold for trip classification. C: Example of trip length as a function of trip number (i.e. temporal order) for a single fly. D: Distribution of log-transformed trip lengths for short (purple) and long (teal) trips for all flies. Due to different normalizations per fly, the two distributions overlap. E: Strip- and boxplot showing trip durations binned according to trip index for short (purple) and long (teal) trips. Boxplots are only shown for bins with a minimum of 8 samples. F: Probability of long trips as a function of trip index. G: Strip- and boxplot showing the normalized cumulative feeding duration at the start of a trip, binned according to trip index. Means per bin are shown as black circles. The bin in which the mean reaches 0.5 is highlighted with shading. H: Scatterplot version of E, with points from short and long trips color-coded. Linear regressions were performed for short and long trips for trips ranges #0-11 and #12-29. Only those regression lines that showed a significant correlation are displayed. The regression results were: short trips (#0-11, solid purple line) R^2^=0.051, *p* = 0.00044851155731283164 *<* 0.001(***); short trips (#12-29, dashed purple line) R^2^=0.004, p=0.4149267399534884 ns.; long trips (#0-11, solid teal line) R^2^=0.0, p=0.937636778156208 ns.; long trips (#12-29, dashed teal line) R^2^=0.015, *p* = 0.08850131135000838 *<* 0.1(*). I: Schematic of various mechanisms that contribute to the scaling of the search pattern. See also Supplemental Figure 6.

### Several behavioral adaptations contribute to the expansion of the search pattern

For the 24 h starved flies foraging for sucrose, short and long trips differed distinctly in their spatial and temporal features. Long trips showed a much larger variance than short trips, and tended to occur later in the trial, interspersed with short trips which flies continued to make throughout their search (**Fig. 5C, E**, see also **Fig. 1F, H**). A closer look at the dynamics of short and long trips revealed that in the early phase of the search (trips 0-5), when flies rarely made long trips, the short trips seemed to gradually increase in size (**Fig. 5E**). Later (trips 6 and onward), the probability of long trips increased, while short trips appeared to remain the same size or slightly decreased (**Fig. 5F**). We next asked whether changes in satiety, approximated by the accumulated feeding time at the start of a trip (**Fig. 5G**), could explain some of these dynamics. We therefore bisected the data into an early phase (up to trip 12), corresponding to the trips when flies on average had not yet reached 50% of their total feeding time, and a late phase (trips 13 and later). We found that short trip lengths increased significantly during the early phase, while long trip lengths increased significantly during the late phase (**Fig. 5H**, see legend for statistics). Despite large differences in feeding dynamics between 24 h and 40 h starved flies, we found similar scaling dynamics for trips performed by 40 h starved flies (**Fig. S6E-H**) suggesting that the scaling effect during the assay was independent of the metabolic state of the fly at the onset of the experiment. Next, we asked if recent feeding, measured by individual visit lengths, could predict subsequent trip durations and vice versa. We found a weak but significant negative correlation between food visits and the subsequent trip length for short but not long trips (**Fig. S6I**, solid lines), that is, long feeding bouts tended to be followed by short trips. Consistent with this, the probability of long trips also decreased with visit duration (**Fig. S6J**). Both observations could simply result from the scaling of short trips in the early search phase and thus reflect the dynamics observed over the course of the whole trial, rather than direct feedback between recent feeding and exploratory trips. More notably, long trips tended to be followed by long feeding bouts (weak positive correlation with the subsequent visit duration, **Fig. S6I**, dashed teal line), which could reflect a drive to maintain energy homeostasis.

Together, our results indicate that two adaptations contribute to the gradual scaling-up of the search pattern: Early on, flies increased the length of their short trips; later, flies increased the probability of taking a long trip and slightly increased the length of these long trips (**Fig. 5I**). These adaptations could be driven by changes in the satiety level during the assay (meals), which in turn is affected by the feeding rate. As the feeding rate is increased by starvation, the metabolic state of the animal at the onset of the meal is another slow-changing key variable defining the foraging strategy of the animal.

### Locomotor adaptations can yield high return rates but do not capture other statistics of behavior

The previous results highlight how flies scale their search behavior over time while still returning to the food spot between trips. We next asked what mechanisms might allow flies to reliably return to the food spot at the end of each exploratory trip, starting with the role of pure locomotor adjustments (Bell, 1985). We segmented behavioral trajectories into runs and turns, approximating straight-line segments separated by circular arcs (**Fig. 6A**, Methods). This allowed us to parameterize each trajectory by its run lengths, turn angles, and turn radii distributions. We fit these distributions separately to the sets of all trips and to only short trips (**Fig. 6B**). We found that the bestfitting parameters of these distributions varied as a function of flies’ distance from the food spot; at farther distances, runs were longer, and turns were broader (**Fig. 6C-D, Fig. S7**). We compared these properties to very short trips with 5 or fewer run and turn segments: Runs were short (**Fig. 6C**, top), and turns were tight (**Fig. 6C**, bottom) and heavily biased in one direction (**Fig. 6D**, bottom). To explore whether these distance-dependent properties were sufficient to enable flies to return to the food spot reliably, we used these distributions to constrain a simple generative model of exploration behavior. When the model was constrained to mimic the properties of very short trips, with short runs and tight, biased turns, it could capture the high return rates (**Fig. 6E**, dark purple lines) and distribution of maximum displacements (**Fig. 6F**, dark purple) observed in the data (grey, all trips). In contrast, when the model was constrained to mimic the properties of all trips, it exhibited much lower return rates and encountered the arena boundary much more often than real flies (**Fig. 6E-F**, light blue/purple). The differences between these models were also reflected in other statistics of behavior; whereas real flies exhibited bimodal distributions of the number of segments and curvilinear distances of their trajectories, corresponding to the long and short trips described above, each generative model was only able to capture one mode (**Fig. 6G-H**): Only the model based on very short trips could capture the short trip mode, whereas both the model fit on short and the one fit on all trips generated trajectories resembling long trips in the data. A look at the simulated trajectories re-sulting from the different models shows that the model based on very short trips generated highly tortuous trajectories that, while resulting in high return rates, did not capture the observed behavior of real flies (**Fig. 6I, Fig. S8**). Together, these results suggest that a generative model of locomotor adaptation can capture the high return rates observed in the data, but it cannot simultaneously capture the behavioral trajectories that real flies use to achieve such high return rates. Thus, flies may adjust their locomotor pattern after food consumption to aid local exploration but are very likely to rely on additional strategies to navigate back to the food from longer distances.

**Figure 6.**
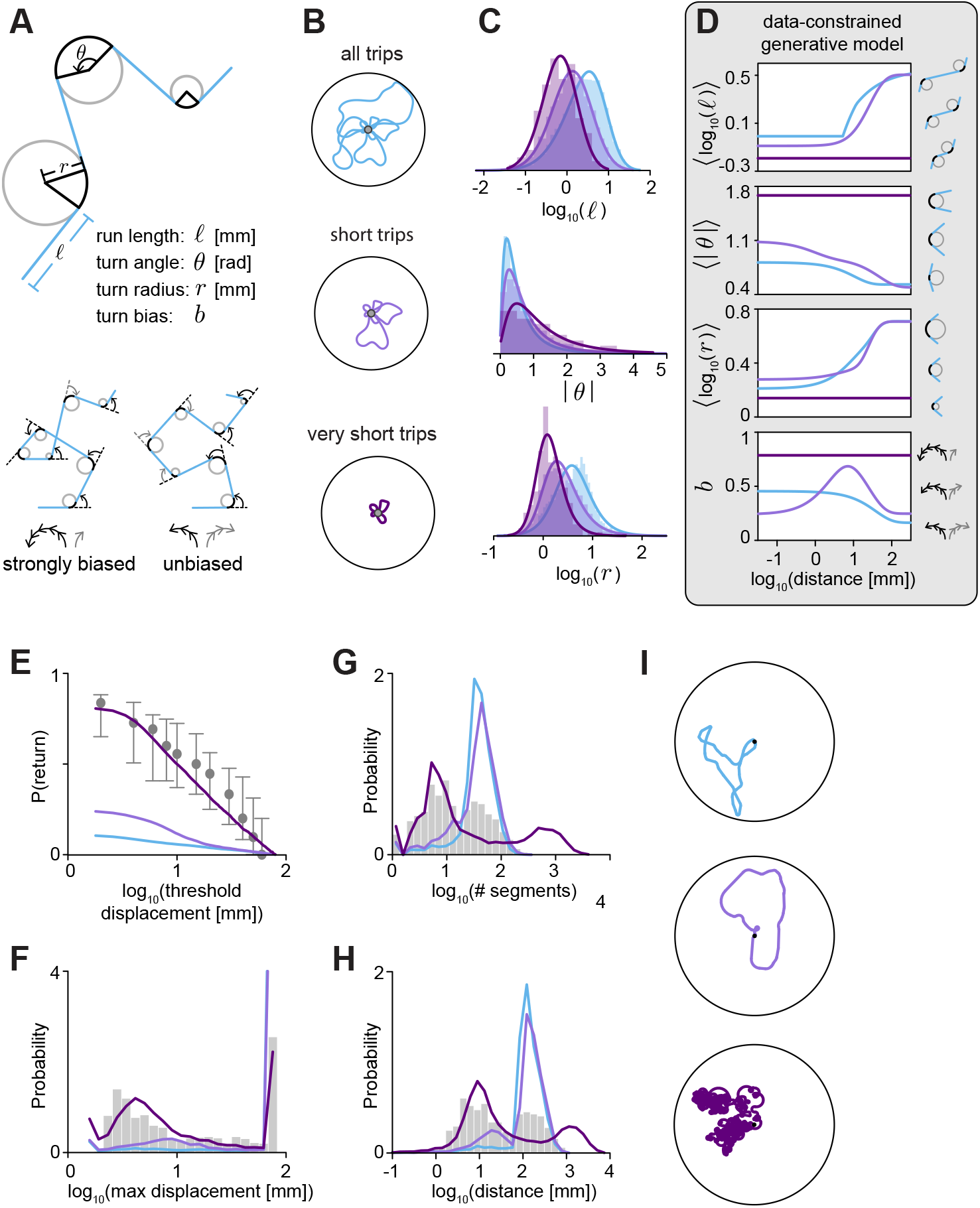
A generative model of locomotor adaptation fails to simultaneously capture high return rates and search trajectory shape. A: Decomposition of trajectories into run and turn segments. Runs are modeled as straight-line segments of length *l*; turns are modeled as circular arcs of radius *r*, angular span *q*, and directional bias *b*. B: Distinction between all trips, short trips, and very short trips. C: Parametric fits to normalized distributions of run lengths (top; SkewNormal), turn angles (middle; LogNormal), and turn radii (lower; SkewLogistic). Fits were performed on data from 24 h starved flies, using all trips (blue), short trips (light purple), and “very short” trips consisting of 5 or fewer segments in total (dark purple). and assumed their statistics stay constant over time. D: We used the parametric functions in panel C to fit distributions of run lengths, turn angles, turn radii, and turn biases for runs and turns taken at different distances from the food spot, illustrated here using the mean of each fitted distribution. We assumed that the statistics of very short trips did not depend on distance. The parameters of these fits form the basis of our generative model (Methods). E-H: Properties of trips that were not used as inputs to the model, compared between data (gray) and different versions of the generative model (color). Properties of the data were computed over all trips; properties of the generative model were computed over 10000 simulated trips. E: Probability of returning to the food spot after exceeding a given displacement. Markers denote median return probabilities computed across flies. Error bars span the 25th to 75th percentile of return probabilities, computed across flies. F-H: Probability distributions of maximum displacement (F), total number of run and turn segments (G), and curvilinear distance along the trajectory (H). Distributions are normalized to sum to one; note that axes exceed a value of one because individual bin widths are much less than one. Both the number of segments and the total distance were computed from the start of a trip to the point when the fly either returned to the food spot or encountered the arena boundary; the large probability mass in the last bin in panel F corresponds to trips that reach the arena boundary. I: Example loops produced by each of the three generative models, shown for loops that reached a maximum displacement of 71 mm (log_10_(*maxdisplacement*) = 1.85). See also Supplemental Figures 7 and 8.

### Humidity sensing appears may guide flies back to the food during long search trips

Because locomotor adjustments alone cannot explain the larger scale search pattern with persistent returns, we concluded that flies searching for food in the dark used additional strategies to return to the food. It has been suggested that flies use path integration to keep track of the food spot location (Behbahani et al., 2021; Corfas et al., 2019; Kim and Dickinson, 2017; Titova et al., 2023). A common assumption is that the insect head direction (HD) system (Seelig and Jayaraman, 2015) is required for path integration (Heinze et al., 2018; Stone et al., 2017). We therefore tested 24 h starved flies with a disrupted HD system by silencing EPG neurons (EPG>Kir and EPG>TNT, Methods) and compared their search behavior to that of wild type (*Canton-S*) flies (**Fig. 7A-E**). We note that other behaviors thought to depend on an intact HD system (menotaxis and visual learning) are impaired when EPG neurons are silenced using the same genetic manipulation we used here (Dan et al., 2024; Giraldo et al., 2018; Green et al., 2019; Haberkern et al., 2022). However, flies with a disrupted HD system showed search patterns that closely resemble those of wild type flies (**Fig. 7B, Fig. S9A**, top) with a comparable number of loops (**Fig. 7C**) and mean loop length (Fig. 7D). The return rates of flies with impaired HD systems also matched those of wild type flies (**Fig. 7E**) and, as we saw in wild type flies, flies with disrupted HD systems scaled their loop distances over time (**Fig. 7B, Fig. S9B**). Thus, perturbing the HD system appeared to have no effect on flies’ ability perform long exploratory trips and re-turn to the food. We therefore considered alternative strategies flies could use to return to the food: sensing odor or humidity gradients associated with the food (**Fig. 7F**). Previous studies suggested that local search does not depend on olfactory cues (Kim and Dickinson, 2017) and we used sucrose presented in agarose drops as a food-source in our experiments specifically to limit olfactory contributions to local search. Nevertheless, we decided to test flies with im-paired olfaction using a odorant receptor co-receptor (*Orco*) knock-out line (Larsson et al., 2004). Motivated by the observation that foraging insects can use humidity cues in addition to olfaction to find food and assess its quality (Harrap et al., 2021; von Arx et al., 2012), we also tested the involvement of hygrosensation in local search. To perturb hygrosensation we used two lines with a disrupted *Tmem63* channel gene (*Tmem63[1]* and *Tmem63[2]*; note *Tmem63* encodes a mechanosensitive ion channel that also plays a role in detecting food texture (Li et al., 2022; Li and Montell, 2021)). *Orco* muta https://github.com/hjmh/foraging-analysis.nt flies showed search patterns that matched searches of wild type flies, consistent with previous findings, while the search pattern of flies with impaired hygrosensation was strongly reduced in scale (**Fig. 7G, Fig. S9A**, bottom row). *Tmem63* mutant flies made fewer and shorter loops compared to wild type and *Orco* mutant flies (**Fig. 7G-I**), suggesting a specific effect on long search loops. Indeed, return rates of flies with impaired hygrosensation were lower for long loops compared to wild type flies (**Fig. 7J**) and long loops were specifically reduced compared to other genotypes tested (**Fig. S9F**). Consistent with the assumption that flies with impaired hygrosensation are deficient in reliably navigating to the food, scaling of the trip lengths was reduced (**Fig. S9C**), and also the time point when *Tmem63* mutants first found the food also showed a large variance (**Fig. S9D**). In contrast to the optogenetic search, however, flies with impaired hygrosensation showed wildtype-like loop tortuosity (**Fig. S9G-H**). Together these observations suggest that hygrotaxis could be potentially used as part of a strategy for returning to the food during local search in the dark.

**Figure 7.**
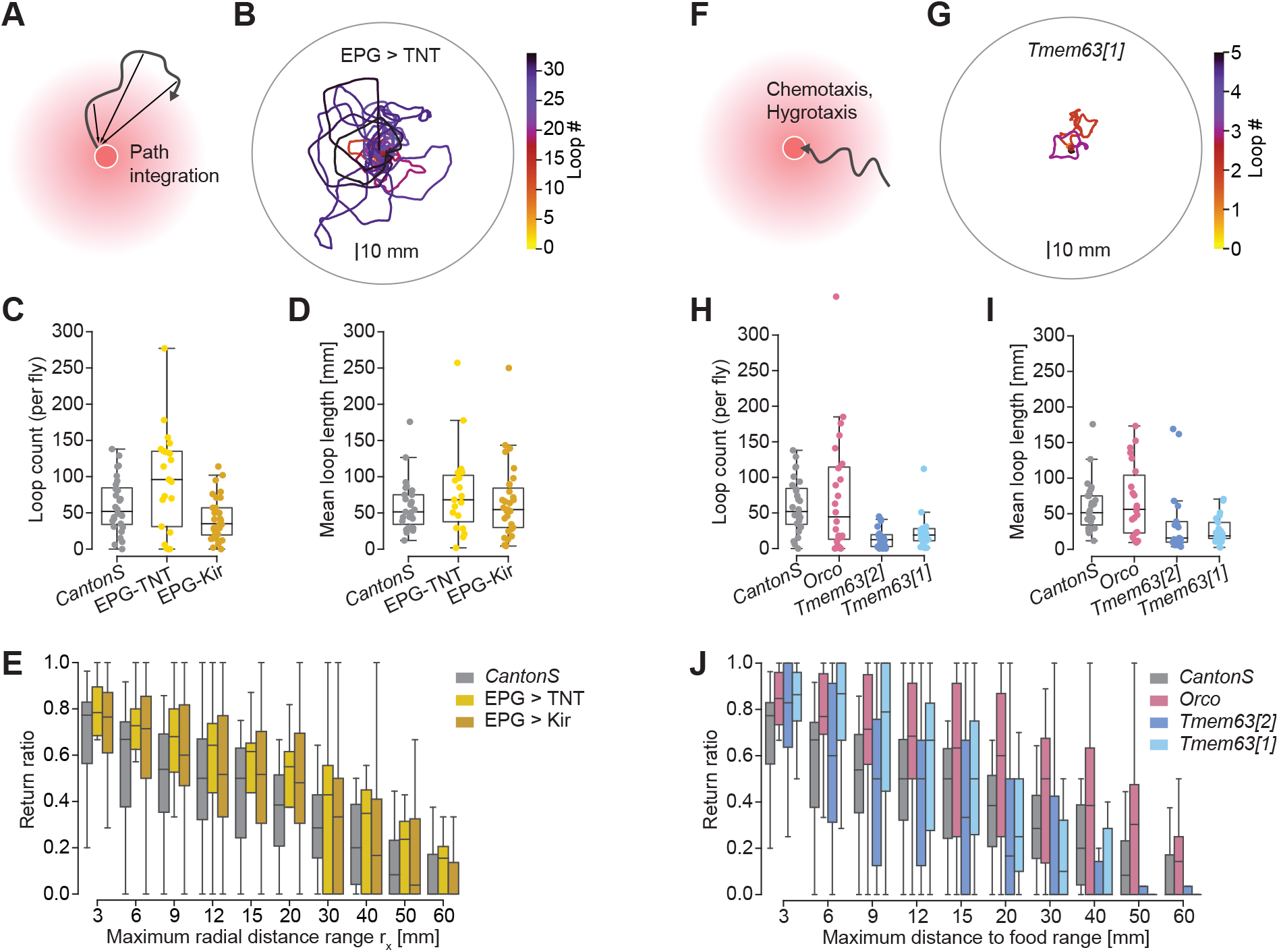
Perturbing individual representational and sensory pathways does not fully disrupt local search. Comparison of local search behavior across different genotypes, all tested with sucrose food after 24 h starvation. Wild type: *Canton-S* (gray, *n* = 27). A-E: Effect of perturbing the head direction system using EPG>TNT (yellow, *n* = 18) and EPG>Kir (ochre, *n* = 30). A: Schematic for how path integration, which requires a head direction estimate, could be used during local search. B: Loops generated by a single EPG>TNT fly, illustrated as in Fig. 1E. C: Boxplot of the total loop count per fly across genotypes. D: Boxplot of the mean loop length per fly across genotypes. E: Return ratio across genotypes (see Fig. 2 and Methods for details). F-J: Effect of perturbing olfaction (*Orco*, rose, *n* = 20) and hygrosensation (*Tmem63[2]*, blue, *n* = 21; *Tmem63[1]*, light blue, *n* = 20), which could help flies find the food through chemo- or hygrotaxis, respectively. F: Schematic of return by chemo- or hygrotaxis. G: Loops generated by a *Tmem63[2]* fly, illustrated as in Fig. 1E. C: Boxplot of the total loop count per fly across genotypes. D: Boxplot of the mean loop length per fly across genotypes. F: Return ratio across genotypes. See also Supplemental Figure 9.

## Discussion

We examined how food quality and internal state modulate the spatiotemporal dynamics of local search in hungry fruit flies and what behavioral strategies flies employ during exploratory search trajectories. Flies’ initial metabolic state and their recent food exploitation determine the size of their exploratory search loops, which increase over time. In early trips, short searches increased in length; in later trips, flies performed more long searches. Flies’ starvation state influenced not just their foraging strategy but also how they re-sponded when they encountered food. The value of this food, sensed via taste and post-ingestive nutritive cues, determined the initiation and duration of search trips. Our analysis of previous studies revealed that searches triggered by optogenetic activation of sugar receptors, which decouples taste and nutritive value, exhibit dynamics that are different from those we observed with actual food emphasizing the importance of metabolic feedback on foraging. We built a simple generative model to assess whether locomotor adaptations were sufficient to produce the returns to food spots that we observed. The simulations suggest that locomotor adaptations can yield high return rates even for long exploratory trips but fail to preserve a low path tortuosity for those long search trajectories. To explore the potential role of other mechanisms for these longer returns, we genetically perturbed different sensory and representational pathways and found that while none of the individual manipulations abolished search, some aspects of the search pattern were altered when perturbing humidity sensing. We propose that the spatiotemporal pattern of local search is shaped by two internal states, a slowly changing starvation-induced metabolic state and a faster changing satiety-related state. The effects of starvation and nutrient deprivation on local search described here recapitulate previous observations in flies and other animals (Münch et al., 2020): depriving flies of nutrients for several days changes behavior for up to three days after refeeding (Ribeiro and Dickson, 2010), leading to increased nutrient-specific feeding (Bell, 1985; Dethier, 1976; Itskov et al., 2014; Ribeiro and Dickson, 2010; Yapici et al., 2016), and biases foraging towards local over global exploration (Bell, 1985; Corrales-Carvajal et al., 2016; Dethier, 1976). In addition to these long-lasting effects of the metabolic state, recent food encounters affect exploratory behavior in flies (Corrales-Carvajal et al., 2016; Goldschmidt et al., 2023; Min et al., 2021; Wang et al., 2020; Wilinski et al., 2019). Our analysis suggests that dynamic adjustments of search behavior related to recent feeding unfold independent of the adjustments of foraging behavior in response to different metabolic states: longer starvation (40 h vs 24 h) led flies to spend more time feeding and biased search to very local exploration around the found food source, while preserving the gradual increase in search radius (Fig. 1). Our analysis of the search patterns induced by activating sugar sensing neurons without providing real food further supports the notion that the expansion of search is driven by satiety signals (Fig. 4), although we note that we do not have a positive control for real-food search in that specific setup. A different study using optogenetic sugar sensor activation in a constraint, one-dimensional assay for local search found evidence for an expanding search pattern (Behbahani et al., 2021). While further studies are necessary to link the observed behavioral dynamics to specific internal states, our data demonstrate that local search behavior emerges from adaptations on multiple time scales, one slow, induced by multi-day deprivation, and a faster one, tracking recent food consumption, that unfolds over minutes. Foraging behaviors including local search in flies are modulated by many internal and external factors such as the metabolic state and sensory signals indicating the presence and quality of food (Munch et al., 2022). Given the importance of successful foraging for the animal’s fitness, it is likely that multiple behavioral strategies exist. Such a degeneracy of behavioral mechanisms could explain the high variability we observed throughout our experiments and complicates the challenge of linking behavior to neural circuits. For example, individual flies chose different exploitative strategies when feeding during local search (Fig. 2C) which could in turn affect the observed dynamics of exploratory behavior as the two are linked through the changes in internal state. Our detailed quantification of local search behavior provides hints for the use of different search strategies and reveals features that help distinguish one strategy from another. Going forward, this will help with connecting the observed behavior to specific neural circuits. Our results suggest that local search in our assay, which takes place in complete darkness, is structured into phases and flies could rely on different exploration strategies during different phases. Exploratory search loops initially span just a few millimeters, but later extend over tens of centimeters. In flies searching around a real food spot path tortuosity increases only slightly with search scale, while flies searching in response to optogenetic activation of sugar sensing neurons take convoluted paths, especially as trips get longer (Corfas et al., 2019; Haberkern et al., 2019) (Fig 4F). Curiously, we saw a search pattern resembling that of optogenetic search emerge from our generative model if we used parameters obtained from matching walking trajectory statistics from time points shortly after flies interacted with the food (Fig. 6). Thus, the very local exploration around a food source in local search as well as optogenetic search could be driven by locomotor adjustments induced by food-related cues such as sweet taste as proposed in early studies of local search (Bell, 1985; Dethier, 1976). Later, during more distal exploration, flies may use alternative strategies that permit efficient return trajectories. The idea that local search is structured into several phases also fits with observation that the gradual increase in search scale was driven by different behavioral mechanisms during early and late search (Fig. 5).

We note that our model does not consider possible correlations between the radius of curvature and the turn angles in the fly data, which may have led us to underestimate the flies’ abilities to return to the food based on locomotor adaptations alone when we matched walking statistics of longer trips. Another simplification is that run and turn statistics across all trips are the same, but these could vary as a function of previous feeding. Thus, our model provides only a lower bound on how well flies may return based on locomotor adaptations alone, leaving the possibility that different search patterns in late stages of local search are due to changes in the locomotor adjustments rather than a change in navigational strategy.

Motivated by the partial mismatch between the trajectories of simulated and real data, we investigated alternative navigational strategies flies could use to return to the food successfully from a distance (Fig. 7). Previous studies suggested that flies rely on path integration (Kim and Dickinson, 2017), but can also follow sensory cues emitted by the food spot or deposited by themselves (Baker et al., 2018; Root et al., 2011; Titova et al., 2023; Tsao et al., 2018; van Breugel, 2021). Our findings suggest that flies can still perform local search in darkness without relying on their head direction system or *Orco*-mediated olfaction. We cannot rule out that flies follow self-deposited olfactory signals or use a path integration-based search strategy interchangeably and that these strategies are difficult to distinguish behaviorally. If it is necessary to disturb multiple pathways at a time to abolish search behavior, a secondary issue arises from potential behavioral differences between flies of different genotypes. Given that perturbation effects were subtle, we cannot rule out that the observed differences in search were due to differences in the genetic background of flies. Still, it is noteworthy that we did see changes in the search pattern in the two mutant lines with impaired humidity sensing. Humidity gradients generated by a food source have not been explored much as guiding foraging in flies. Further investigation will be necessary to confirm that indeed the humidity sensing function tied to the TMEM63 channel or its role in mechanosensation was responsible for the observed behavioral effects. Given the importance of finding and returning to food sources for an animal, the ability to use several different strategies during local search may be an adaptation to achieve robustness to perturbations and changing conditions. This is in line with observations in other insects that use a wide variety of sensory cues, including hygrosensation, to localize and identify flowers (Raguso, 2004; Rands et al., 2023). The temporal increase in scale is a key feature of local search behavior in flies can be seen as a strategy to dynamically balance exploitation and exploration as suggested before (Corrales-Carvajal et al., 2016). A pattern of loops that gradually expand in scale has also been described in other contexts, for example nest searches (Degen et al., 2016; Schultheiss and Cheng, 2011) and learning walks or flights (Woodgate et al., 2016; Zeil and Fleischmann, 2019) in hymenopterans. In the case of learning walks and flights, the insect performs this behavior around a site of interest, typically the nest, to which it needs to return later. The increase in search scale is thought to reflect a strategy to successively collect more information, that is explore the environment while remaining oriented using local cues (Bertrand and Sonntag, 2023). Notably learning flights can also be observed in the context of foraging around flowers, even in non-central place foragers (Robert et al., 2018). Thus, both local search behavior and learning flights and walks balance exploration with exploitation and their algorithmic similarity raises the possibility that these two behavioral patterns may share neural mechanisms. In conclusion, our study highlights the inherent dynamics in the local search behavior of foraging animals: multiple feedback loops encompassing both off-the-food exploration and onthe-food exploitation, as well as interactions between behavioral actions and internally generated, latent state variables representing satiety and metabolic state. We show that the observed temporal and spatial patterns in local search behavior emerge from these feedback dynamics with flies expressing different behavioral patterns and possibly switching strate-gies throughout their search. The intricate interplay of feedback processes makes it challenging to establish direct causal relationships between sensory signals, internal state and neural computations. Our findings, suggesting a subdivision of local search into different stages that could rely on different internal and external signals, therefore provide an important step toward linking foraging behavior to neural circuits.

## ACKNOWLEDGEMENTS

We thank Célia Baltazar for technical assistance with the nutrient spot experiments, C.C., Claire Managan and Gudrun Ihrke for support through Project Technical Resources, and the Janelia Visiting Scientist program for support of the collaboration. We thank members of the Ribeiro, Jayaraman and Hermundstad labs for discussions and feedback on early versions of the manuscript. S.S.C. was supported by the Janelia Undergraduate Scholars Program. D.G. was supported by a doctoral fellowship PD/BD/114273/2016 from the Portuguese Foundation for Science and Technology (FCT). The project leading to these results has received funding from the grant PTDC/MED-NEU/4001/2021 from the Portuguese Foundation for Science and Technology (FCT) awarded to C.R. Research at the Centre for the Unknown is supported by the Champalimaud Foundation. Research at Janelia was funded by the Howard Hughes Medical Institute.

## Methods

### RESOURCE AVAILABILITY

#### Lead contact

Further information and requests for resources and reagents should be directed to and will be fulfilled by the lead contact, Hannah Haberkern.

#### Materials availability

This study did not generate new unique reagents.

#### Data and code availability

- All datasets generated in this study will be made available upon publication on data repositories. We will provide raw (movies, metadata files) and preprocessed data. The corresponding accession numbers are listed in the key resources table.
- All original code is available through the Github repositories listed in the key resource table.
- Any additional information required to reanalyze the data reported in this paper is available from the lead contact upon request.

### EXPERIMENTAL MODEL AND STUDY PARTICIPANT DETAILS

Our study involved experiments performed in two locations, at Janelia Research Campus in Ashburn VA (abbreviated as JR below) and Champalimaud Research Center (CR below). Differences in fly rearing are detailed below. At JR, flies were reared in an incubator set to 23° C temperature, 60% relative humidity and 16:8 light:dark cycle. Fly food was prepared according to the following recipe: agar 0.7%, cornmeal 8.0%, inactivated yeast 2.0%, molasses 5.0%, propionic acid 0.50% and tegosept 0.20%. Agar was brought to boil in a kettle. Cornmeal and dry yeast were mixed together in water before adding to the boiling agar. Molasses was added and the mixture was boiled for 10 minutes on average. Tegosept and propionic acid were added after turning off the heat. For starvation experiments, flies were moved to a vial containing 1% agarose (Sigma-Aldrich Co. LLC, St. Louis, MO, USA) and a moist Wattman filter paper 24 h or 40 h prior to introducing them to the local search assay. At CR, flies were kept at 25°C, 70% relative humidity in a 12 h/ 12 h light-dark cycle and reared on a yeast-based medium (per liter of water: 8 g agar (NZYTech, Portugal), 80 g barley malt syrup (Próvida, Portugal), 22 g sugar beet syrup (Grafschafter, Germany), 80 g cornflour (Próvida, Portugal), 10 g soya flour (A. Centazi, Portugal), 18 g instant yeast (Safinstant, Lesaffre, France), 8 mL propionic acid (Argos), and 12 mL nipagin (Tegosept, Dutscher, France) (15% in 96% ethanol) supplemented with instant yeast granules on the surface (Saf-instant, Lesaffre, France)).

Datasets with the following laboratory stocks were used: *Canton-S* MH (Jayaraman Lab stock, derived from Martin Heisenberg stock), *Canton-S* (Ribeiro Lab stock), *Orco* mutants (olfactory receptor mutant as a control for the effect of food smell), SS96 (split Gal4 labeling EPG neurons), UAS-TNT in *Canton-S* MH background, UAS-Kir2.1 in *Canton-S* background, and *Tmem63[1]* and *Tmem63[2]* humidity sensory mutants. Complete genotypes are listed in the key resources table. Only female flies were used in experiments.

### METHOD DETAILS

Behavioral experiments in this study were performed in two locations, at Janelia Research Campus in Ashburn VA (abbreviated as JR below) and Champalimaud Research Center (CR below). The experimental setups used in the two locations were similar with some differences that we highlight below. Data from both locations was later fed into a shared analysis pipeline.

#### Local search assay

The local search assay was modeled after 32. We recorded walking traces of individual female flies for 45 min while they explored a large, dark arena with a single food source placed at the center. The food was presented in the form of a small (5 µl) drop of 1% agarose which was either prepared with a sucrose solution (125 mM, sucrose from Sigma-Aldrich Co. LLC, St. Louis, MO, USA) or water (0 mM). The arena was 170 mm in diameter and about 3 mm high with the edges cut in a small tooth-pattern to prevent flies from walking along the edge. The arena was covered with a glass plate that was treated with Sigmacote® (Ref. SL2, Sigma-Aldrich Co. LLC, St. Louis, MO, USA) during experiments to prevent flies from leaving the arena and from walking on the ceiling. For the JR assay, the arena was placed inside an incubator that ensured stable temperature and humidity (25° C temperature and 50% relative humidity) as well as complete darkness. Prior to the experiment, the floor of the arena was wiped with a moist tissue paper and the agarose drop was gently placed at its center. Flies were then transferred from the starvation vial to the boundary of the arena with a mouth aspirator. For the CR assays, the dimensions of the arena were identical to the ones used for JR assays. We placed a 5 µl spot of 100 mM sucrose, sucralose or sorbitol with 1% agarose into the arena center using a multipipette. We coated the arena lids (black, IR-transmissive acrylic glass) with 20 µL Sigmacote® (Ref. SL2, Sigma-Aldrich Co. LLC, St. Louis, MO, USA) to prevent flies from leaving the arena and from walking on the ceiling. The black acrylic glass ensured that flies performed the assay practically in darkness. The CR assays were performed a temperature- and humidity-controlled behavioral chamber (25° C, 70% relative humidity).

#### Image-based tracking setups

For JR datasets, videos were recorded with BIAS (IORodeo, Pasadena, CA) in .ufmf format using a FLIR Grasshopper3 camera (GS3-U3-23S6M, Teledyne FLIR LLC, USA) at 24 Hz at a resolution of 1920 px × 1200 px resulting in a pixel resolution of 5.1 px per mm. Walking trajectories were subsequently extracted from the raw video data using Ctrax (Branson et al., 2009). For CR datasets, we used a high-throughput tracking setup that can record four flies in individual arenas using a custombuilt acrylic arena holder26. Arenas were placed above an IR backlight for video recording using a FLIR Flea3 camera (FL3-U3-32S2M-CS, Teledyne FLIR LLC, USA) acquiring videos at 30 Hz at a resolution of 1408 px × 1408 px. This resolution resulted in an average real-world pixel resolution of 7.8 px per mm. We used a custom, real-time tracking algorithm created in the Bonsai framework (Lopes et al., 2015). Fly centroids and trajectories were extracted using a background subtraction method, and for each recorded frame we measured the x and y-position of the centroid, major and minor axis as well as the angle of the centroid. We also developed custom-written Python scripts to determine the head position of the fly using an offline algorithm. This algorithm compares the pixel intensity of both extreme points of the centroid along the major axis to determine the head, which is darker than the tail position due to the transparency of the wings. We applied a proximity rule (Gomez-Marin et al., 2011), in which the position closer to the previous head position is assigned as head position, to propagate the head detection through time. For jumps this rule can become invalid, so we detected jumps as segments to re-apply the pixel intensity comparison. Furthermore, we validated correct head position assignment, by comparing the movement vector to the heading vector, which should largely align because flies mostly walk forwards. The accuracy of the head detection by manual visual inspection scored more than 98% of frames (1000 frames randomly sampled).

#### Experimental conditions

We tested how starvation state and food quality (pure agarose vs. agarose with 125 mM sucrose, 100 mM sucrose vs. 100 mM sucralose vs. 100 mM sorbitol) affected search behavior in this assay. To control the starvation state, we wet starved flies for 24 h or 40 h in vials with pure agarose instead of regular fly food. In some additional experiments, we deprived flies of specific nutrients (i.e., from carbohydrates and amino acids) based on a holidic diet described previously (Leitao-Gonçalves et al., 2017) using the exome-matched formulation with food preservatives (Piper et al., 2017). See below for a full list of experimental conditions and dataset sizes.

**Table 1.** Overview of experimental conditions and sample sizes.

| Genotype | Starvation | Food (agarose based) | Collection location | Sample size | Figures |
| --- | --- | --- | --- | --- | --- |
| *Canton-S* (MH) | 24 h | sucrose, 125 mM | Janelia | 28 | 1, 2, 4-7, SI1, SI2, SI4-9 |
| *Canton-S* (MH) | 40 h | sucrose, 125 mM | Janelia | 29 | 1, 2, 4, SI1, SI2, SI5, SI6 |
| *Canton-S* (MH) | 24 h | water | Janelia | 26 | SI4, SI5 |
| *Canton-S* | 3d -SAA | sucrose, 100 mM | Champalimaud | 14 | 3, SI3 |
| *Canton-S* | 3d -SAA | sucralose, 100 mM | Champalimaud | 14 | 3, SI3 |
| *Canton-S* | 3d -SAA | sorbitol, 100 mM | Champalimaud | 14 | 3, SI3 |
| *Orco* | 24 h | sucrose, 125 mM | Janelia | 30 | 7, SI9 |
| SS96 >TNT (*Canton-S* MH) | 24 h | sucrose, 125 mM | Janelia | 21 | 7, SI9 |
| SS96 >Kir | 24 h | sucrose, 125 mM | Janelia | 31 | 7, SI9 |
| *Tmem63*[2] | 24 h | sucrose, 125 mM | Janelia | 23 | 7, SI9 |
| *Tmem63*[1] | 24 h | sucrose, 125 mM | Janelia | 21 | 7, SI9 |

### QUANTIFICATION AND STATISTICAL ANALYSIS

#### Preprocessing pipeline

In this study, we used data collected in two different locations (see Local search assay) and we designed a preprocessing pipeline that allowed us to represent the collected data in a shared format. This enabled direct comparisons across datasets. The code for the preprocessing pipeline can be found here, alongside jupyter notebooks demonstrating the use: https://github.com/degoldschmidt/foraging-datapreprocessing. Briefly, the preprocessing consists of three steps: First, metadata about the arena and food spot were generated. For JR datasets, metadata about arena size and food position were extracted from the .ufmf recordings by extracting the first frame of the recording in fiji using a custom macro (getFirstUFMFframe_batch.ijm) and subsequent processing in python (generate_ctrax_metadata.py). For CR datasets, custom-written Python code which is part of the tracking analysis (described above) has been used to automatically detect and extract metadata information, such as arena size and food position, described in Goldschmidt et al. (2023). Finally, metadata about the fly’s genotype, experimental conditions and arena size were saved into a standardized metadata file in yaml format. In a second step, the Ctrax and Bonsai output, which contained basic trajectory information, such as the fly position, body axis and time steps, was converted into a pandas data frame with the following standardized variables:

- *frame*: index of video frame number
- *dt*: time interval between current and next frame [s]
- *body_x*: x coordinate of body position (centroid) [px]
- *body_y*: y coordinate of body position (centroid) [px]
- *head_x*: x coordinate of head position (centroid) [px]
- *head_y*: y coordinate of head position (centroid) [px]
- *major*: length of major axis of centroid [px]
- *minor*: length of minor axis of centroid [px]
- *angle*: estimated body angle [rad]

In the third step, derived trajectory quantities such estimated feeding, frame-by-frame labels for ethogram states (see below) and path segments (see below) were computed:

- *body_speed*: translational speed of the body position [mm/s] smoothed by
- *head_speed*: translational speed of the head position [mm/s].
- *angular_speed*: angular speed [rad/s]
- *distance_center*: distance to arena center [mm]
- *distance_patch_0*: distance to the food spot [mm]
- *ethogram*: classification of discrete behaviors based on kinematic parameters, e.g., angular and body speed (see below for detailed description).
- *segment*: broader classification of local search segments (see below for detailed description).

For a given dataset, all trajectory per-frame variables were saved in the form of a single, concatenated pandas data frame and stored as a feather file, using the label-parameters ‘fly’, ‘condition’, ‘genotype’ to separate experiments within the file.

#### Ethogram classification of trajectories

We applied thresholds onto the time series of translational as well as rotational speeds to segment the trajectory on a frame-by-frame basis into 5 ethogram states:

- Border: Frames during which the fly was close to the arena border using a Schmitt trigger (76 mm and 79 mm for the large arena, 22.5 mm and 25 mm for the small arena respectively).
- Turn: Turns were detected as frames with filtered rotational velocity above a 120 deg/s threshold. The turn start was defined as the initial crossing of this threshold and the turn end as the frame at which rotational velocity dropped below this threshold.
- Peaks in angular speed and speed difference between head and body (findTurns function)
- Run: Frames are labeled as runs based on filtered body speed using a Schmitt trigger (low threshold = 1.0 mm/s, high threshold = 3.0 mm/s).
- Feeding: Feeding events can override turns and are based on the fly’s proximity to the food spot and low head speed (frames that are not labeled as runs).
- Other: Any remaining frames are labeled as “other”.

#### Classification of trajectory segments

We further classified trajectories on a larger scale into different walking segments. This classification is based partly on the ethogram classification described above.

- Border: Defined as described under ethogram classification.
- Visit: Visits are all consecutive feeding frames that do not exceed a leaving threshold from the spot.
- Loop: Loops are path segments that are flanked by visit segments.
- Arrival: Arrival segments start at the border and end at the food (are followed by a visit).
- Departure: Departure segments start at the food (follow a visit) and end at the border (or with the end of the trial).
- Miss: These are segments that start and end at the border (or with the end of the trial)

#### Analysis of feeding dynamics

To analyze feeding dynamics, we analyzed a behavioral proxy for detecting feeding micromovements (Corrales-Carvajal et al., 2016; Goldschmidt et al., 2023). Segments of feeding micromovements in which the fly resides at the patch without leaving beyond a defined distance threshold were classified as feeding visits. The statistics of feeding visit durations, i.e., how long a fly stays feeding, was characterized by the average of all visit durations. The frequency of feeding visits was quantified by the total number of visits.

#### Analysis of loop and trip structure

We analyzed higherorder features of the trajectories by quantifying various statistics on a per-loop or per-trip basis. Here loops were defined as described above (see “Classification of trajectory segments”) and trips were defined as a segment, in which the fly leaves the food spot and explores the arena irrespective of contacts with the border before returning to the food spot. For each loop or trip, respectively, we computed which trip/loop it was for an individual fly, whether it was a long or short trip/loop, the duration [s], the path length [mm] and the maximum radial distance [mm] reached. We also computed the duration of the feeding visit before [s], the duration of the feeding visit after the trip/loop [s], the total cumulative feeding time [s] up to the trip/loop and the cumulative feeding normalized by the total duration of feeding (ranging from 0 to 1). These parameters were then combined in pandas DataFrames together with several descriptive variables such as the experimental condition and the fly id. Data from multiple of these data frames was combined for analyses that compared across genotypes. Statistical tests and regressions were performed using the scipy.stats package. Throughout the paper the sample size refers to the number of flies. The code for analyzing loop and trip structure can be found here: https://github.com/hjmh/foraging-analysis.

#### Modeling of locomotor adaptations

We used our generative model to simulate the trajectories of model flies. Each simulated trajectory was initialized at the center of a circular region with a radius of 1.5 mm, which represented the food spot in the experiment. For each trial, the initial heading angle of the model fly was randomly drawn from the range (-π/2, π/2) relative to the normal vector at the exit point on the boundary of the food spot.

In our model, we assume trajectories are composed of alternating straight run segments and curved turn segments; we model these curved segments as circular arcs. Based on analysis of the very first segment of trips in the data, we assume that each trajectory has an equal probability of starting with a run or turn segment. The length of run segments *l*, the absolute turn angles |Θ|, and the radius of curvature *r* of the turn segments are each random variables drawn from the following probability distributions inferred from the data: log_10_ *l ∼ SkewNormal*, |Θ| *∼ LogNormal, log*_10_*r ∼ Type*1*GeneralizedLogistic*.

A trajectory ends when it either returns to the food spot or when it encounters the arena boundary. To account for the fly body length, if the displacement from the boundary of the food spot during a trajectory exceeds a value *ϵ* = 1 mm (approximately half the body length of a fly), we consider the fly to have returned to the food spot when its displacement from the food spot boundary next falls below *ϵ*.

#### Statistical feature extraction for modeling

Since our model assumes that trajectories only contain alternating run and turn segments, we first removed other types of behavior (grooming, etc.) from all trips before analyzing the remaining run and turn segments. For each run segment, we calculate the total distance traveled. For each turn segment, we extract the total distance, the overall change in heading angle from the start to the end of that segment, as well as the turn direction (clockwise or anti-clockwise). Assuming that the turn segments can be approximated as circular arcs, the effective radius of curvature of each turn segment is then given by the (total distance traveled during the segment)/|turn angle|.

To choose the most suitable distribution for each of the random variables (run lengths, turn angles and radius of curvature), we used the python Fitter package to fit a range of standard distributions (to either the data or log10(data)) and compared their fit.

All chosen distributions are parameterized by 3 variables. When using data from all trips and from all short trips, we binned the runs and turns based on distance traveled since leaving the food spot at the start of the segment (bin edges are chosen such that there are an equal number of samples in each bin), and we fitted the distributions separately for each of these bins. This allows us to obtain values for the parameters of the fitted distributions at discrete time points. We then fit either a power law function, an exponential function, or a sum of two exponentials (to allow for non-monotonic changes) to get a function for how each of the parameters change as a function of distance. If there is no clear trend that the parameter changes with distance, we assume that the value of that parameter is fixed over time.

When using data from very short trips (trips with 5 or less segments; these make up 40% of all short trips), the time dependence is not extracted because the trip distances tend to be very short. We therefore use the fitted distributions from all runs or turn segments in these trips and in the simulations, assume that they remain the same throughout the trip. We note that run and turn statistics extracted from all trips or all short trips were not weighted by trip length. Thus, a single trip among all short trips that is much longer than most of the short trips, would contribute much more to the statistics of the runs and turns.

When inferring the probability of turning in a preferred direction *p*_*bias*_, we first determine for each trip the preferred turn direction by counting the number of turns in the clockwise and anticlockwise directions and choosing the direction with more turns. Based on the turn direction for each segment, we then determine if it is in the preferred direction. The values of *p*_*bias*_ are then obtained by averaging across all the relevant trips across all flies in the same starvation condition.

## Supplementary material

**A. Supplemental Information.** Supplemental Figures 1-9 with figure legends.

