## Supplemental information for "Exploration and exploitation are flexibly balanced during local search in flies"

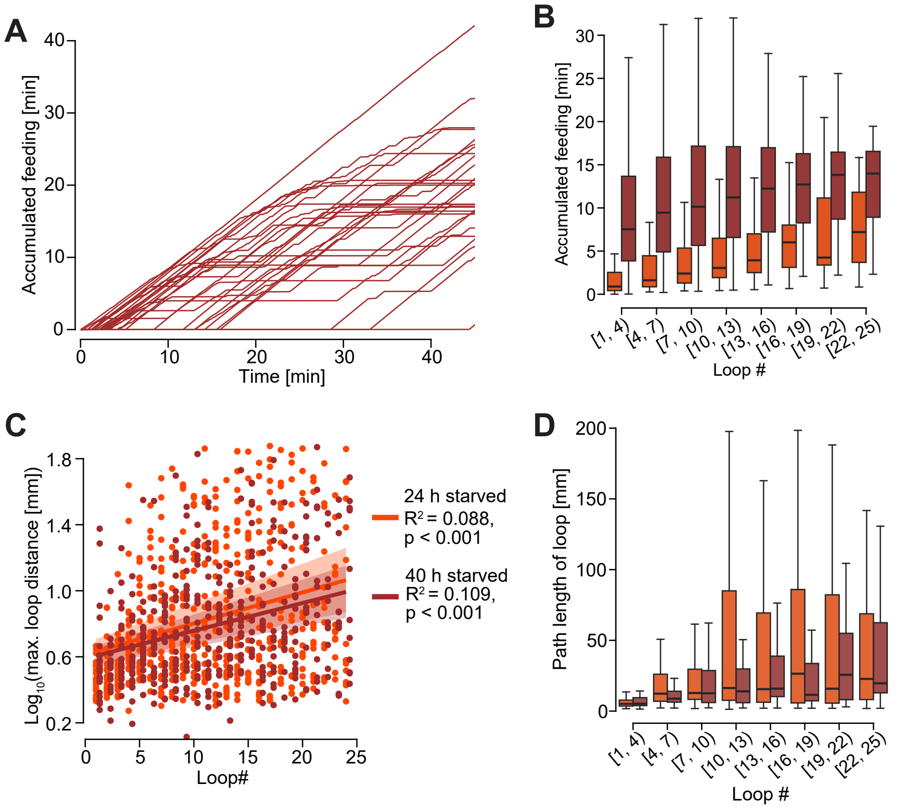


### Supplemental Figure 1

**A**: Accumulated feeding for 40 h starved wild type flies, visualized as in Figure 1D.

**B**: Boxplot of the accumulated feeding at the start of a loop, visualized as in 1H. Experimental groups are color coded: 24 h starved flies in orange (n=28) and 40 h starved flies in dark red (n=29).

**C**: Scatterplot of loop number vs the logarithm of the maximum radial loop distance. The logarithmic transform was chosen due to the long-tailed distribution of loop distances. The two experimental groups are color coded as in B. Regression lines were generated using ordinary least squares linear regression (24 h starved wild type: $R^2$=0.088, p=6.270106396335541$e^{-12}$; 40 h starved wild type: $R^2$=0.109, p=9.417834233499424$e^{-14}$). Shaded region indicates 95% confidence interval.

**D**: Boxplot of loop path lengths with loops binned according to their temporal order, comparing the 24 h and 40 h starved flies.

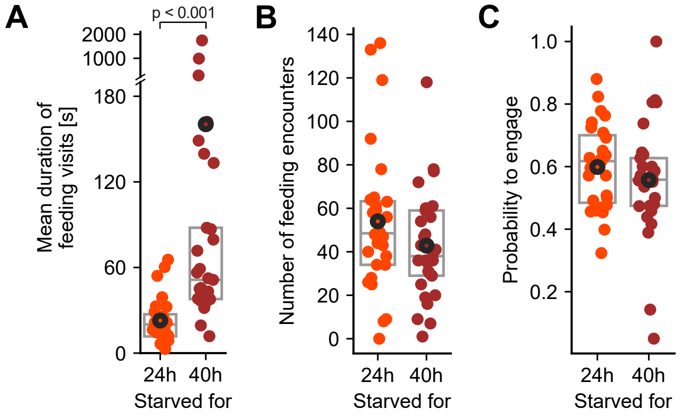


### Supplemental Figure 2

**A**:  Mean duration of feeding visits in seconds (s) for 24 h and 40 h starved flies.

**B**: Number of feeding visits for 24 h and 40 h starved flies.

**C**: Number of food spot encounters for 24 h and 40 h starved flies.

**D**: Probability to engage with a food spot for 24 h and 40 h starved flies. The probability is defined as the ratio of number of feeding visits divided by the number of food spot encounters.

**A-D**: Data points are calculated for individual flies. Boxes indicate the median and interquartile range (IQR, i.e., between 25th and 75th percentile), and circle markers indicate the mean. P-values are obtained by performing Wilcoxon rank-sum tests, n = 28-29.

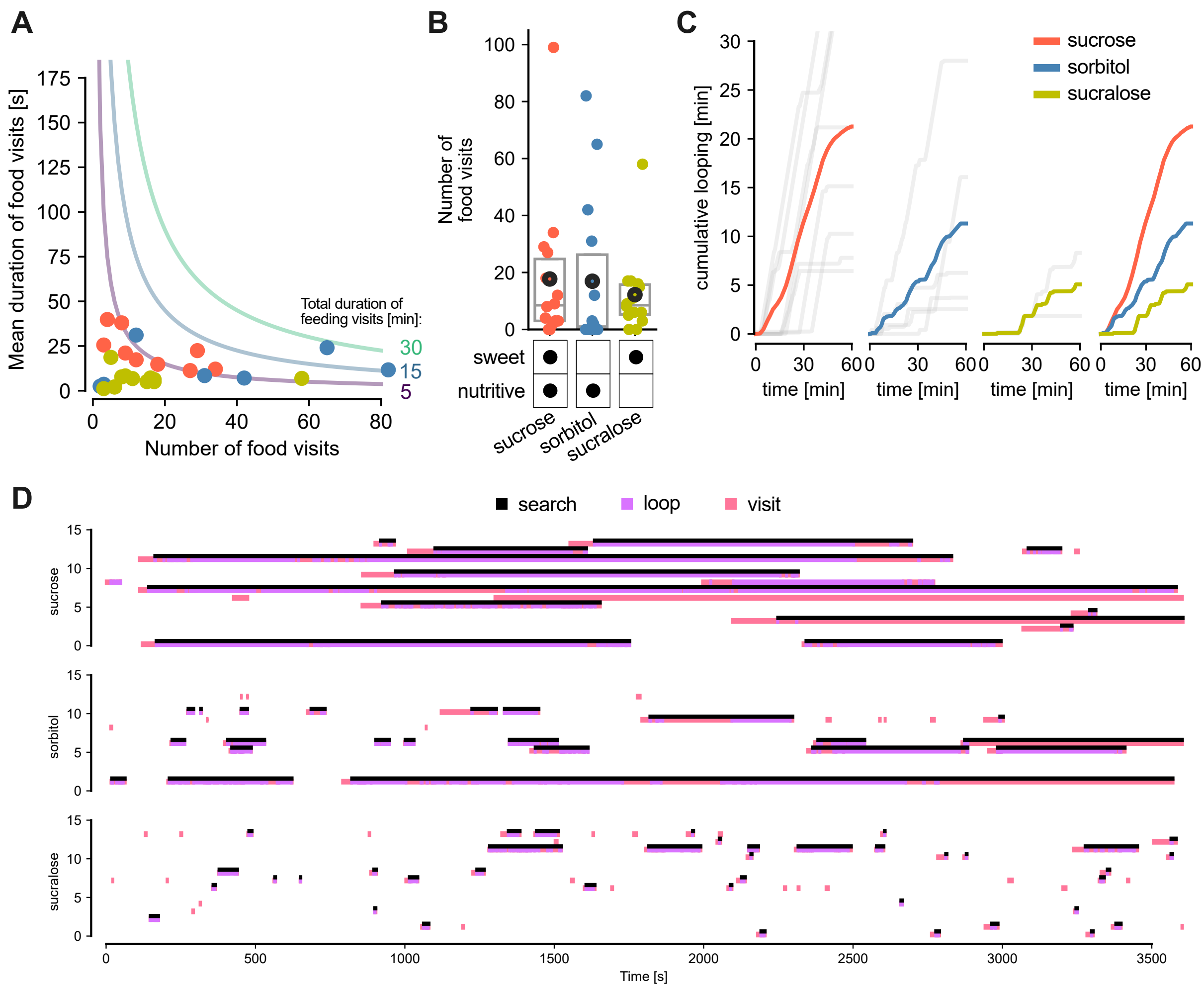


### Supplemental Figure 3

**A**: Mean duration of feeding visits in seconds (s) for deprived flies presented with 100 mM sucrose, 100 mM sorbitol or 100 mM sucralose.
**B**: Number of feeding visits for deprived flies presented with 100 mM sucrose, 100 mM sorbitol or 100 mM sucralose.
**A-B**. Data points are calculated for individual flies. Boxes indicate the median and interquartile range (IQR, i.e., between 25th and 75th percentile), and circle markers indicate the mean. P-values are obtained by performing Wilcoxon rank-sum tests, n =14.
**C**: Cumulative loops in minutes over the course of the trial for each fly which performed loops (gray lines) as well as the average across flies presented with 100 mM sucrose (red line), 100 mM sorbitol (blue line) or 100 mM sucralose (olive green line).
**D**: Ethogram for individual flies searching for sucrose (top), sorbitol (middle), and sucralose (bottom), which perform feeding visits (purple rectangles) and loops (pink rectangles) over time. We defined search bouts as the segments when a fly leaves a food patch and performs alternating loops and feeding visits before eventually departing to the border.

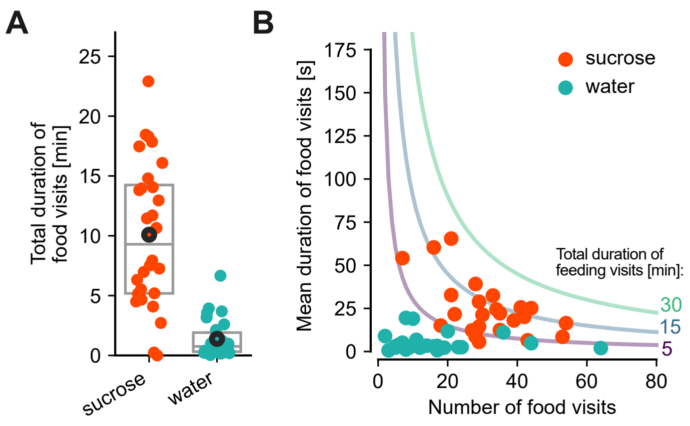


### Supplemental Figure 4

**A**: Total duration of feeding visits in minutes (min) for 24 h starved flies presented with a sucrose (red circles) or water (cyan circles) spot.

**B**: To reach different total durations of feeding visits (purple to green lines), flies may modulate how often to visit (x-axis: number of visits) and/or how long to visit (y-axis: mean duration of feeding visits in min). Data points are calculated for individual, 24 h starved flies presented with sucrose (red circles) or water (cyan circles), respectively.

**C**: Mean duration of feeding visits in seconds (s) for 24 h starved flies presented with a sucrose or water spot.

**D**: Number of feeding visits for 24 h starved flies presented with a sucrose or water spot. **A, C-D**. Data points are calculated for individual flies. Boxes indicate the median and interquartile range (IQR, i.e., between 25th and 75th percentile), and circle markers indicate the mean. P-values are obtained by performing Wilcoxon rank-sum tests, n = 28.

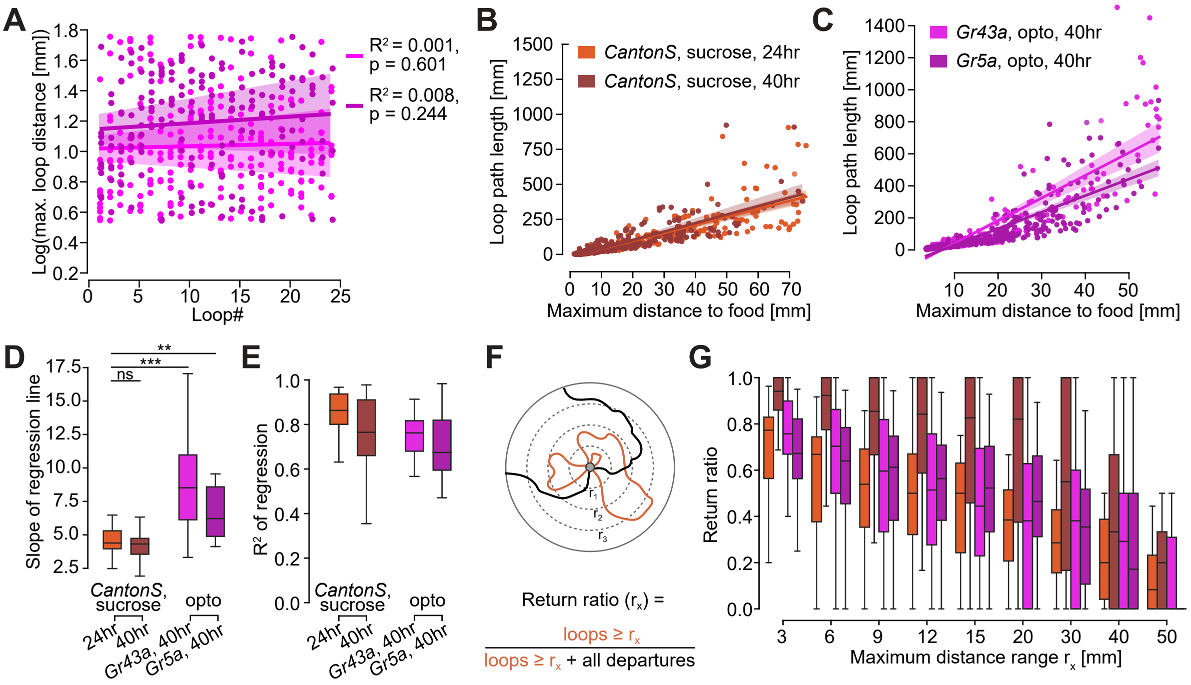


### Supplemental Figure 5

**A**: Scatterplot of loop number vs the logarithm of the maximum radial loop distance. The two experimental groups are color coded as in C. Regression results are shown as described in Fig. 4C. Full parameters: Gr43a>CsChrimson: $R^2$=0.001, p=0.6005009258188643; Gr5a>CsChrimson: $R^2$=0.008, p=0.24412469659864977;

**B**: Scatterplot of maximal distance to food vs path length for loops for 24 h and 40 h starved flies. The regression line was calculated over all data points within the two experimental groups. Shaded region shows 95% confidence interval based on bootstrapping.

**C**: Same as B, but for the two experimental groups with optogenetic stimulation.

**D, E**: Results from a linear regression on the maximum loop distance vs path length calculated per fly.

**D**: Slope of the regression line. T-tests were performed to test for differences between the 24 h starved wild type group and the three other experimental groups (no difference with 40 h starved wild type: t=-0.12006798851250716, p=0.9049304534656346, df=48.0; Gr43a>Chrimson: t=-4.487227118877648, p=6.789309811957789e-05, df=37.0; Gr5a>Chrimson: t=-3.1296048432747314, p=0.003408127168079529, df=37.0)

**E**: R2 value of the regression.

**F**: Definition of the return ratio: the number of loops divided by the sum of loops and departures, as a function of the maximum distance from the food that is reached during loops.

**G**: Return ratio as a function of radial distance from the food spot. Experimental groups are color coded as in B, C.

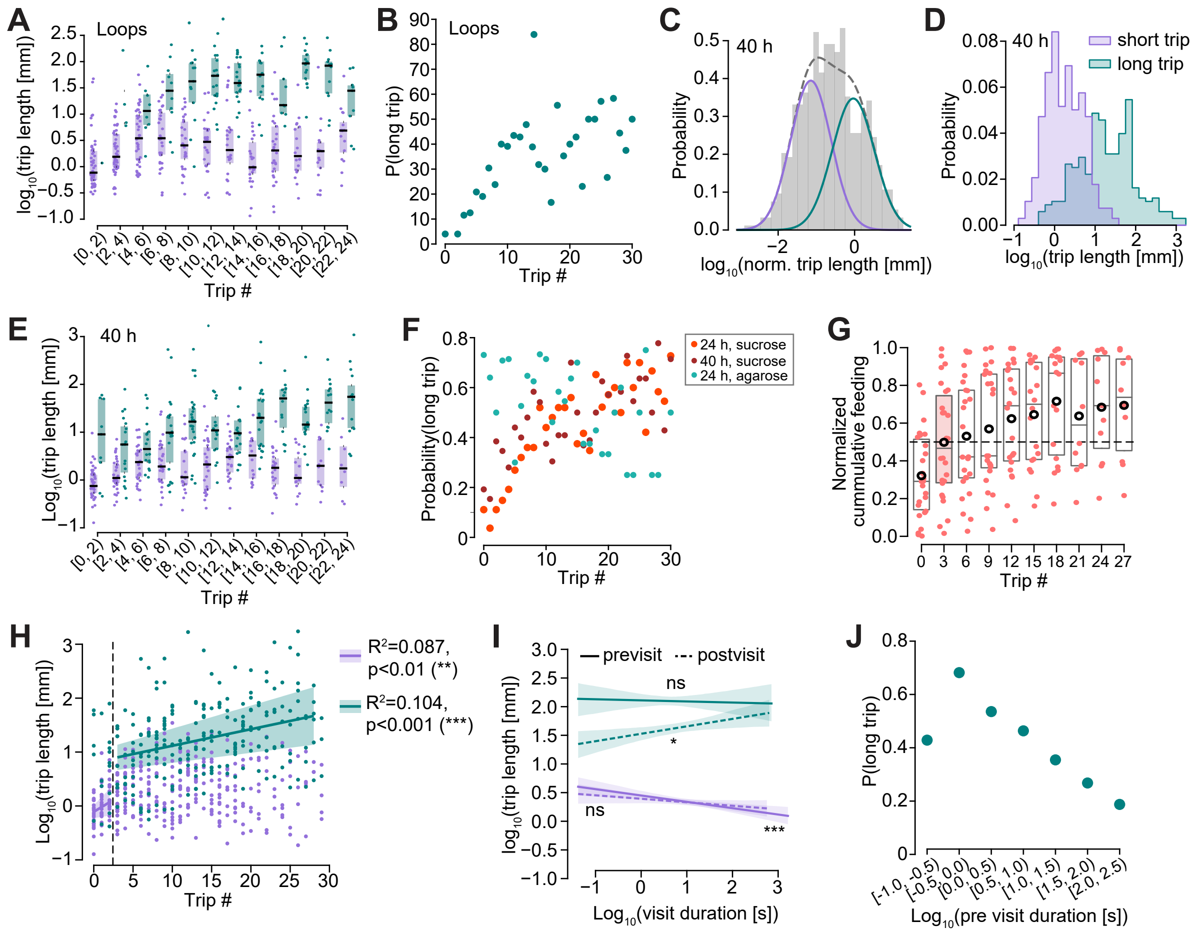


### Supplemental Figure 6

**A**: Strip- and boxplot as in Fig. 5E but only for loops (not all trips).
**B**: Probability of long trips as a function of trip index but only considering loops.
**C**: Bimodal distribution of log-transformed, normalized trip lengths (gray shaded histogram, see methods for details) visualized as in Fig. 5B but for 40 h starved flies. Parameters of the gaussians: (short trips, purple, mean = -1.15, sd = 0.53, weight = 0.52) and long trips (teal, mean = -0.03, sd = 0.55, weight = 0.48).
**D**: Distribution of log-transformed trip lengths for short (purple) and long (teal) trips for all flies as in Fig. 5D but for 40 h starved flies.
**E**: Strip- and boxplot as in Fig. 5E but for 40 h starved flies.
**F**: Probability of long trips as a function of trip index for three experimental groups: 24 h and 40 h starved with sucrose food, 24 h starved with pure agarose food.
**G**: Strip- and boxplot of normalized feeding durations visualized as in Fig. 5G but for 40 h starved flies.
**H**: Scatterplot version of E, with points from short and long trips color-coded. Linear regressions were performed for short and long trips for trips ranges #0-3 and #4-29. Only those regression lines that showed a significant correlation are displayed. The regression results were: short trips (#0-3, solid purple line) $R^2$=0.087, p=0.0084844544823124 < 0.01 (**); short trips (#4-29, dashed purple line) $R^2$=0.0, p=0.9630380788178914 ns.; long trips (#0-3, solid teal line) $R^2$=0.07, p=0.6809130048833728 ns.; long trips (#4-29, dashed teal line) $R^2$=0.104, p=1.0703189152087837e-06 < 0.001 (***).
**I**: Linear regression plot of the log-transformed trip lengths as a function of log-transformed visit durations, with regression against pre-trip visits in solid lines and post-trip visits in dashed lines. The data was split for short (purple) and long (teal) trips. Regression results: pre-visit, short trip: $R^2$=0.025, p=0.0007184686350419591 (***); pre-visit, long trip: $R^2$=0.0, p=0.7668570370655754 (ns.); post-visit, short trip: $R^2$=0.006, p=0.10037555381782735 (ns.); post-visit, long trip: $R^2$=0.026, p=0.022278734876886964 (*).
**J**: Probability of long trips as a function of the log-transformed, previous visit duration.

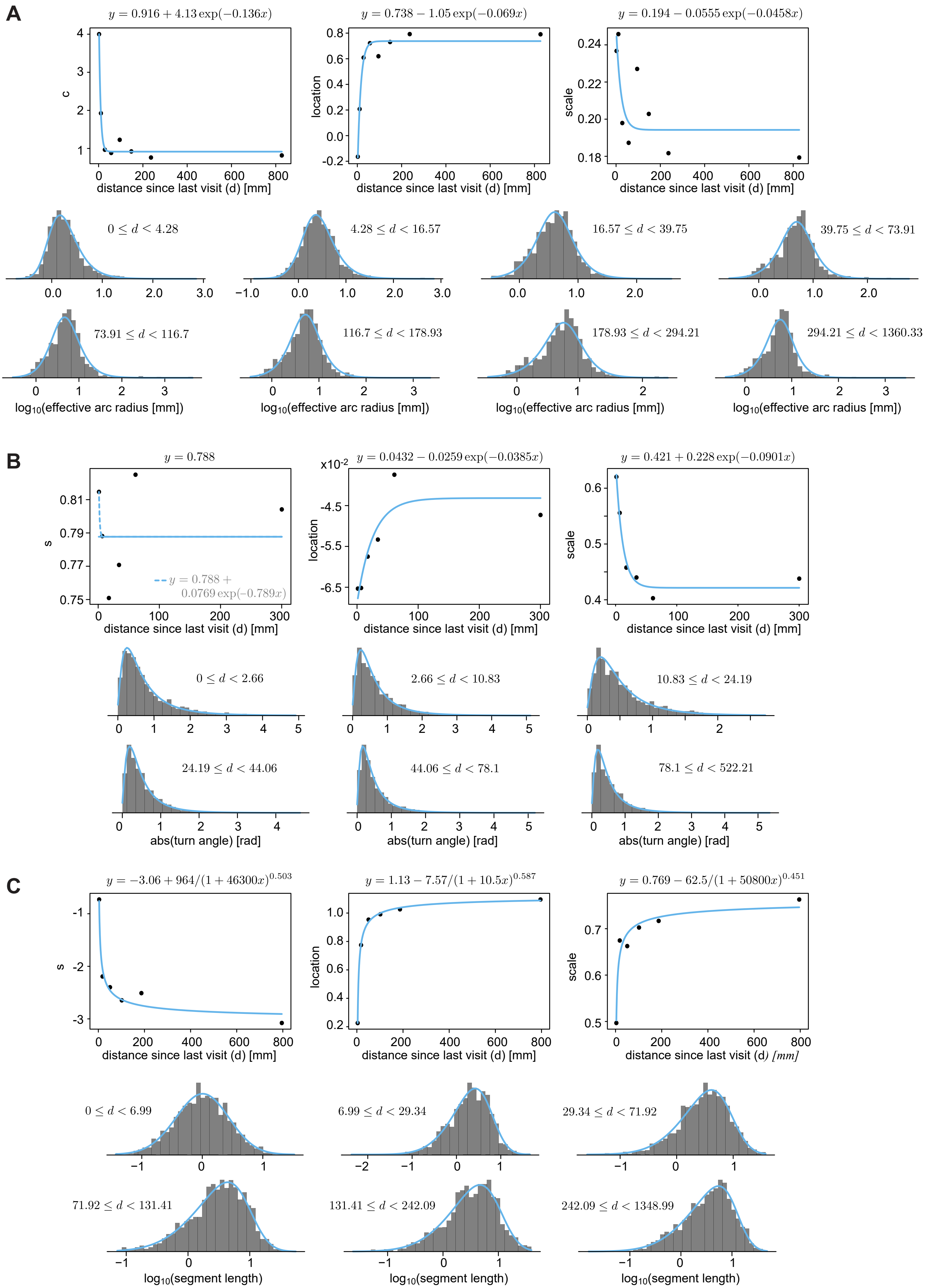
A
$y = 0.916 + 4.13\exp(-0.136x)$
c
distance since last visit (d) [mm]
$y = 0.738 - 1.05\exp(-0.069x)$
location
distance since last visit (d) [mm]
$y = 0.194 - 0.0555\exp(-0.0458x)$
scale
distance since last visit (d) [mm]
$0 \le d < 4.28$
$4.28 \le d < 16.57$
$16.57 \le d < 39.75$
$39.75 \le d < 73.91$
$73.91 \le d < 116.7$
$116.7 \le d < 178.93$
$178.93 \le d < 294.21$
$294.21 \le d < 1360.33$
$\log_{10}$(effective arc radius [mm])
B
$y = 0.788$
s
$y = 0.788 + 0.0769\exp(-0.789x)$
distance since last visit (d) [mm]
$y = 0.0432 - 0.0259\exp(-0.0385x)$
x10$^{-2}$
location
distance since last visit (d) [mm]
$y = 0.421 + 0.228\exp(-0.0901x)$
scale
distance since last visit (d) [mm]
$0 \le d < 2.66$
$2.66 \le d < 10.83$
$10.83 \le d < 24.19$
$24.19 \le d < 44.06$
$44.06 \le d < 78.1$
$78.1 \le d < 522.21$
abs(turn angle) [rad]
C
$y = -3.06 + 964/(1 + 46300x)^{0.503}$
s
distance since last visit (d) [mm]
$y = 1.13 - 7.57/(1 + 10.5x)^{0.587}$
location
distance since last visit (d) [mm]
$y = 0.769 - 62.5/(1 + 50800x)^{0.451}$
scale
distance since last visit (d) [mm]
$0 \le d < 6.99$
$6.99 \le d < 29.34$
$29.34 \le d < 71.92$
$71.92 \le d < 131.41$
$131.41 \le d < 242.09$
$242.09 \le d < 1348.99$
$\log_{10}$(segment length)

#### Supplemental Figure 7:

How parameters of fitted distributions vary with distance since last visit, shown for **(A)** SkewLogistic fit to log10(turn radii), **(B)** LogNormal fit to absolute turn angles, and **(C)** SkewNormal fit to log10(run lengths). Fits of distributions were performed on data from 24 h starved flies, using all run/turn segments within specified distance range across all trips. The variation of parameters with distance were fitted to either an exponential **(A, B)** or power law function **(C)**. For the shape parameters in the left panel of **(B)**, there is no clear trend that the parameter changes with distance (dashed line). We therefore approximate this parameter to be fixed at a value given by the plateau of the fitted exponential function (solid line).

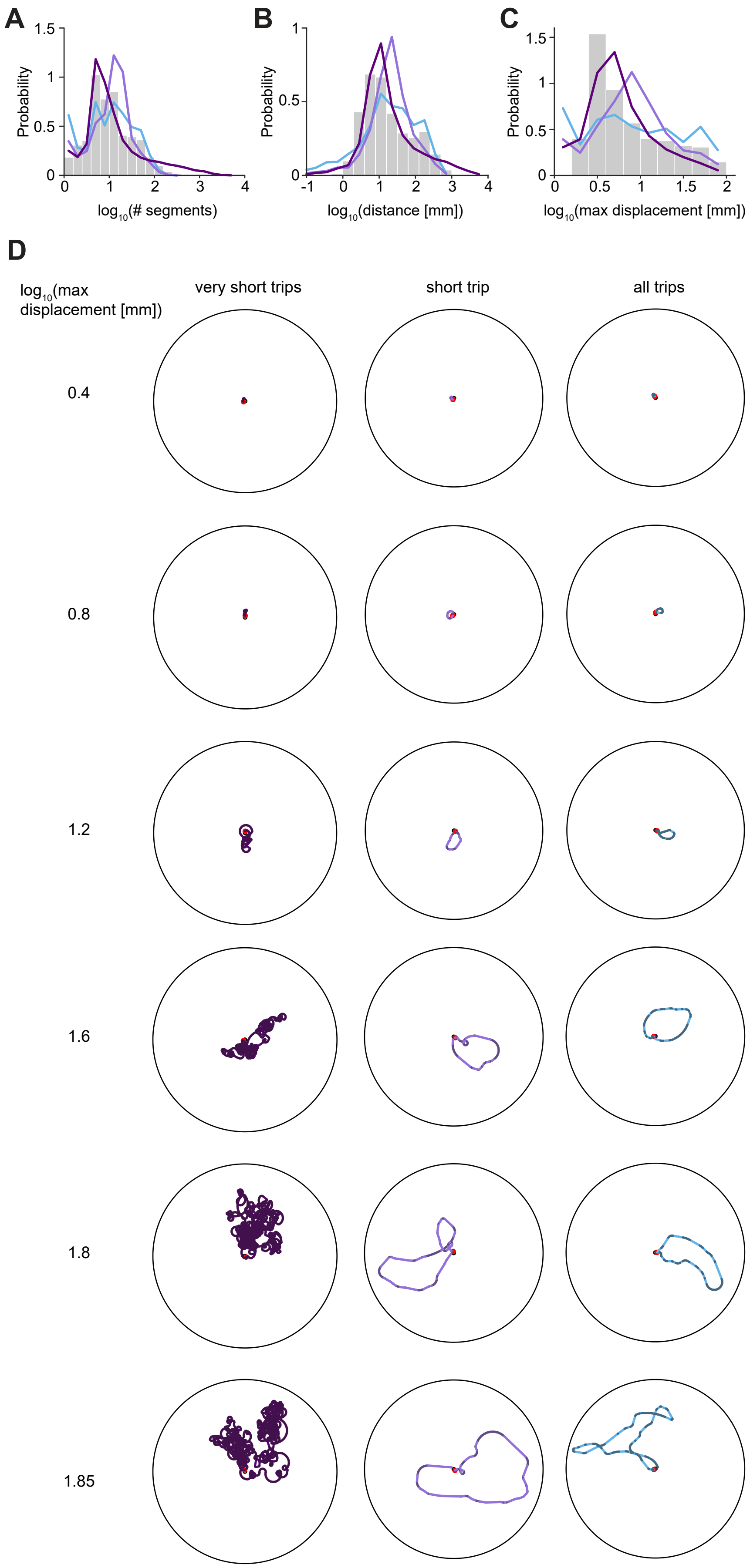


**Supplemental Figure 8:**

**A-C**: Same as Fig. 6F-H, but shown for loops, instead of trips.

**D**: Same as Fig. 6I but segmented into runs and turns and shown for a range of maximum displacements.

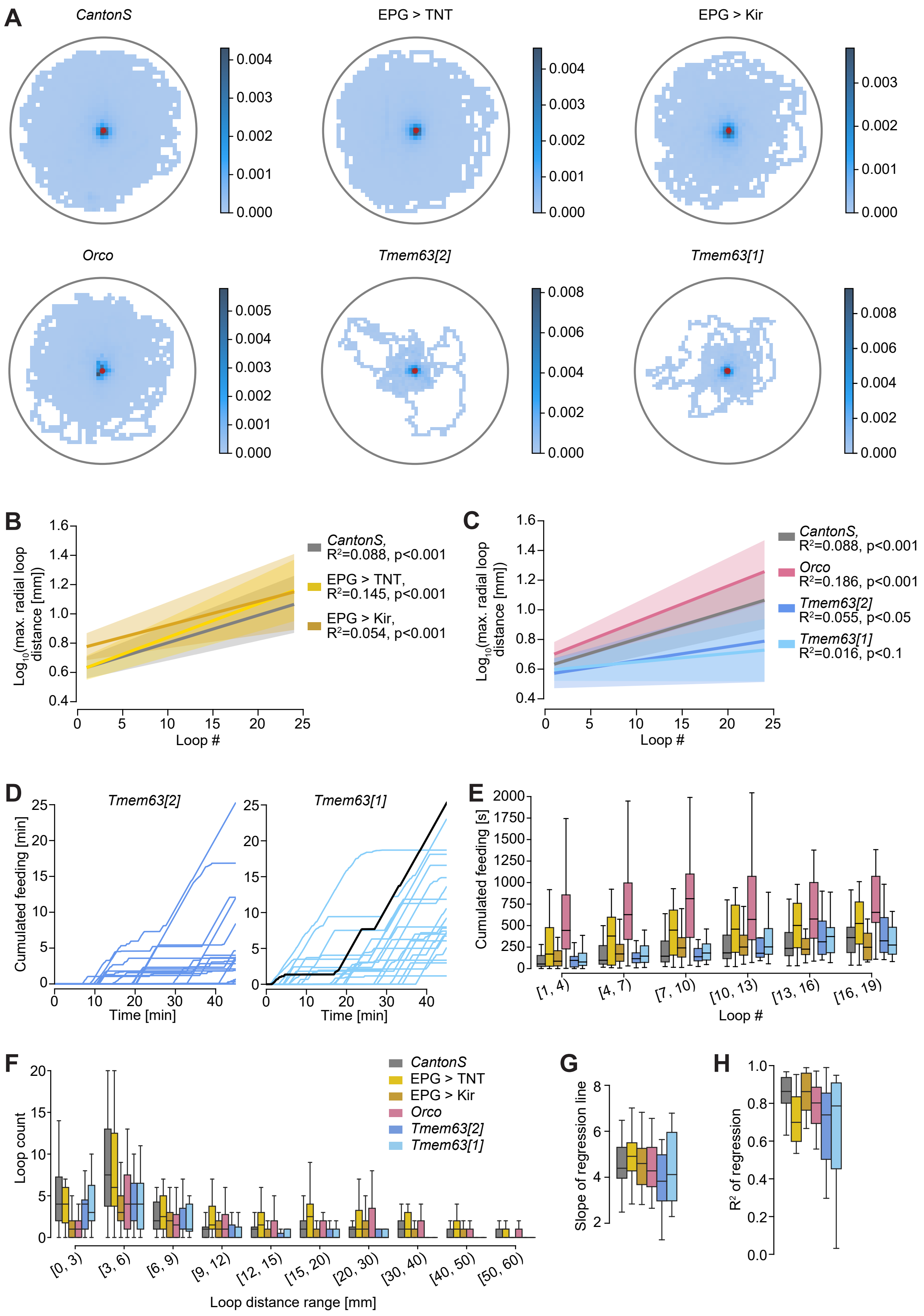

A
CantonS
EPG > TNT
EPG > Kir
Orco
Tmem63[2]
Tmem63[1]
B
Log10(max. radial loop distance [mm])
Loop #
CantonS, R²=0.088, p<0.001
EPG > TNT, R²=0.145, p<0.001
EPG > Kir, R²=0.054, p<0.001
C
CantonS, R²=0.088, p<0.001
Orco R²=0.186, p<0.001
Tmem63[2] R²=0.055, p<0.05
Tmem63[1] R²=0.016, p<0.1
D
Tmem63[2]
Tmem63[1]
Cumulated feeding [min]
Time [min]
E
Cumulated feeding [s]
Loop #
[1, 4)
[4, 7)
[7, 10)
[10, 13)
[13, 16)
[16, 19)
F
Loop count
CantonS
EPG > TNT
EPG > Kir
Orco
Tmem63[2]
Tmem63[1]
[0, 3)
[3, 6)
[6, 9)
[9, 12)
[12, 15)
[15, 20)
[20, 30)
[30, 40)
[40, 50)
[50, 60)
Loop distance range [mm]
G
Slope of regression line
H
R² of regression

### Supplemental Figure 9:

**A**: Two-dimensional search density measured as area normalized residency during loops and departures for the 6 tested genotypes. *Canton-S* (WT, n=27), silenced compass neurons (EPG > TNT, n=18; EPG >Kir, n=30), impaired olfaction (*Orco*, n=20), impaired humidity sensing (*Tmem63[2],* n=21; *Tmem63[1],* n=20).

**B**: Results of ordinary least squares linear regression of the log(maximal radial loop distance [mm]) as a function of the loop number. The log transform was chosen to account for the long-tailed distribution of loop distances. Groups are color coded as in 7C-E. The lines indicate the regression line, the shaded regions the 95% confidence intervals. Full parameters: *Canton-S*: $R^2$=0.088, p=6.270106396335541$e^{-12}$; EPG>TNT: $R^2$=0.145, p=5.00441609387856$e^{-15}$; EPG>Kir $R^2$=0.054, p=1.4944867565848888$e^{-5}$.

**C**: Same as B, but for *Orco* and *Tmem63* mutants. Full parameters: *Canton-S*: $R^2$=0.088, p=6.270106396335541$e^{-12}$ (same as above) *Orco*: $R^2$=0.186, p=1.956135775264042$e^{-15}$; *Tmem63[2]* $R^2$=0.055, p=0.01550402066871149; *Tmem63[1]* $R^2$=0.016, p=0.05717230412321111.

**D**: Cumulative feeding duration as a function of trial time for individual *Tmem63* mutant flies. The time series corresponding to the fly shown in Fig. 6G is highlighted in black.

**E**: Boxplot of cumulative feeding times reached at the start of a loop as a function of the loop number for different genotypes (see F for color code).

**F**: Boxplot of loop counts per fly, binned according to the maximal distance from the food per loop. Data from different genotypes is color-coded as indicated in the legend.

**G, H**: Results of a linear regression of the loop path length to loop distance ratio. Regressions were performed per fly and resulting slopes (**G**) and $R^2$-values (**H**) are visualized as boxplots. See F for color-code.
